# Advancing Nuclei Isolation from Frozen Human Heart for Single-Nucleus RNA Sequencing Applications

**DOI:** 10.64898/2026.03.19.712944

**Authors:** Rocco Caliandro, Ruggero Belluomo, Jermo Hanemaaijer-van der Veer, Roelof-Jan Oostra, Maurice J.B. van den Hoff, Reinier A. Boon, Monika M. Gladka

## Abstract

While single-cell RNA sequencing (scRNA-seq) has been the first widely adopted single-cell transcriptomic approach, its reliance on fresh tissue samples has substantially limited its applicability to clinically relevant specimen. Single-nucleus RNA sequencing (snRNA-seq) overcomes this constrain by enabling transcriptomic profiling from frozen material. However, isolating high-quality nuclei from frozen cardiac tissue remains technically challenging due to the dense extracellular matrix, complex tissue architecture, and heterogeneous cellular composition of the heart. To address these challenges, numerous nuclei isolation protocols have been adapted and optimized, resulting in substantial methodological heterogeneity across studies. Despite the widespread use of snRNA-seq in cardiac research, a robust and standardized nuclei isolation protocol that consistently yields high-quality nuclei from frozen human heart tissue is still lacking. Here, we present a comprehensive, end-to-end protocol for nuclei isolation from frozen human left ventricle, along with a detailed downstream pipeline for snRNA-seq data analysis. Our hybrid nuclei isolation strategy integrates multiple sequential clean-up steps designed to preserve nuclear integrity and RNA quality prior to sequencing. Compared with commonly used nuclei isolation protocols, this approach yields substantially higher number of nuclei while maintaining comparable numbers of detected genes and counts, even at lower sequencing depth. Adoption of this protocol may reduce technical variability across studies and facilitate more reproducible snRNA-seq analyses of human cardiac tissue.

## INTRODUCTION

Single-cell-based transcriptomic technologies have advanced our understanding of tissue complexity by dissecting gene expression at the level of individual cells or nuclei.^1^ Unlike bulk RNA sequencing, which averages signals across the heterogeneous cell populations, these approaches preserve cell-to-cell variability and allow the identification of transcriptional programs within specific cellular subpopulations that may be critical for disease development and progression.^1,2^ In cardiovascular research, such resolution has proven particularly valuable, as subtle changes within cardiomyocyte, fibroblast, or immune cell subsets can have profound effects on cardiac function and pathology.^1^ Single-cell RNA sequencing (scRNA-seq) has been widely used in mapping cardiac cell populations and characterizing disease-associated transcriptional signatures. However, the requirement for fresh tissue limits its applicability, especially in human studies where access to freshly collected cardiac material is often limited.^3^ Single-nucleus RNA sequencing (snRNA-seq) overcomes this limitation by enabling transcriptomic profiling from frozen tissue, thereby providing access to archived samples, large biobank collections, and rare patient material.^3^ As a result, snRNA-seq has become an essential tool for studying human cardiac biology and disease. Despite its advantages, snRNA-seq from frozen human heart tissue presents substantial technical challenges, particularly during nuclei isolation. Nuclei isolation protocols must be tailored to the unique structural and biochemical properties of different tissues. Therefore, methods optimized for other organs cannot simply be transferred to the heart. The adult human heart is characterized by dense extracellular matrix,^4^ high mitochondrial content,^5^ and large, fragile cardiomyocytes, features that complicate efficient nuclei recovery and increase the risk of RNA degradation or contamination.^1^ To address these technical challenges, multiple nuclei isolation protocols have been adapted and applied in cardiovascular research. However, a systematic comparison of the available protocols is still lacking, leaving a methodological heterogeneity that hinders the standardization of nuclei isolation and snRNA-seq in cardiac tissue. Studies that performed snRNA-seq on frozen human cardiac tissue vary widely in reported sequencing depth, nuclei yield, and per-nucleus resolution. Many protocol-focused studies emphasize technical feasibility and cell type diversity, while providing limited information on post-sequencing quality metrics such as Unique Molecular Identifier (UMI) counts per nucleus, gene counts per nucleus, or percentage of mitochondrial reads detected. Similarly, large-scale atlas studies often prioritize the number of nuclei profiled, with less emphasis on detailed reporting of per-nucleus quality metrics. In the absence of standardized benchmarks and systematic evaluation of post-sequencing metrics, it remains difficult to objectively assess the performance of different nuclei isolation approaches.^1^ To address this gap, we compiled post-sequencing quality metrics from published studies that performed snRNA-seq on nuclei isolated from frozen human left ventricle (LV) and compared with data generated using our in-house optimized nuclei isolation protocol. This comparison demonstrates that our protocol yields higher nuclei recovery and a comparable number of genes and UMI detected per nucleus for low depth of sequencing, while maintaining low levels of mitochondrial RNA contamination. Here, we provide a standardized and reproducible framework for nuclei isolation from frozen adult human heart tissue for snRNA-seq.

## RESULTS

Across published studies that employed nuclei isolation from frozen human left ventricular tissue, substantial methodological variability can be observed, reflecting the technical challenges of handling the adult human myocardium. Most protocols rely on detergent-based lysis, using ionic or non-ionic detergents such as Triton X-100, NP-40, Tween-20, or CHAPS, in combination with mechanical disruption, such as Dounce homogenization or tissue pulverization (**Table 1**). However, downstream clean-up strategies to remove debris, mitochondria, and ambient RNA differ considerably. Some studies apply only post-lysis filtration (one-step clean-up) while most studies employ two-step clean-up strategies, combining filtration post-lysis with either density gradient (sucrose/iodixanol) or fluorescence-activated cell sorting (FACS) (**Table 1**). The choice of clean-up strategy depends largely on the tissue features (e.g., high collagen content) and the intended downstream application. Each of these technical variations in the nuclei isolation method could affect the quality of the isolated nuclei, potentially improving or compromising quality and depth of the downstream sequencing. However, a proper assessment requires comparing post-sequencing quality metrics across methods.^1^ Hence, we retrieved curated snRNA-seq datasets from studies that performed 10x Genomics 3’ snRNA-seq sequencing in human LV samples and compared post-sequencing quality metrics across studies. For each dataset, we determined the median number of nuclei recovered per sample (n_Nuclei), median UMI counts per nucleus (n_Counts), and median number of detected genes per nucleus (n_Genes), and the median percentage of mitochondrial reads per nucleus (pct_Counts_mt), after removal of low-quality nuclei and empty droplets. For comparability, only LV samples were included, and all available samples per study were pooled, irrespective of experimental condition of belonging, or age and sex of the donors. When specified by the authors, we also included sequencing specifications like the number of nuclei per reaction of the 10x Chromium Controller or the targeted per-nucleus sequencing depth. This analysis was restricted to studies that made curated snRNA-seq datasets publicly available. In **Table 2**, we summarized n_Nuclei, n_Counts, n_Genes, and pct_Counts_mt per study, while distributions of these post-sequencing quality metrics are shown in **Supplementary Figure 1A-D**. Although no universal cut-offs exist to unequivocally discriminate between low- and high-quality snRNA-seq datasets,^6^ commonly used thresholds provide guidance for assessing the robustness of an experiment. A dataset containing fewer than ∼ 5000 nuclei is generally underpowered for detecting less-abundant cell populations and for robust differential expression analysis. Among the studies that performed snRNA-seq from LV we analyzed, only three out of seven studies^7–9^ met this criterion (**Table 2**, **Supplementary Figure 1A**), suggesting limited nuclei recovery likely due to sub-optimal nuclei isolation strategies rather than insufficient loading. In contrast, most studies achieved acceptable transcript recovery, with n_Counts ≥ 2000 UMIs per nucleus and pct_Counts_mt ≤ 3% (**Table 2**, **Supplementary Figure 1B-C**).^7–13^ However, per-nucleus gene detection remained modest. Most datasets reported 1000-1500 genes per nucleus for a targeted depth of sequencing of 20000-50000 reads per nucleus. ^7,8,10–13^ Only the study by *Youness et al., 2025,*^9^ achieved substantially higher gene count of around 2700 genes per nucleus (**Table 2**, **Supplementary Figure 1D**), with a targeted sequencing depth of 80000 reads per nucleus. While ∼ 1000-1500 genes per nucleus is sufficient to detect major transcriptional changes, this range represents the lower end of what can be achievable using the 10x platform. Studies performing snRNA-seq on frozen human brain tissue, for instance, have reported up to ∼ 6500 genes per nucleus (with a median depth of sequencing of 30000-40000 reads per nucleus),^14^ indicating that current cardiac nuclei isolation methods remain suboptimal and often yield low-powered snRNA-seq experiments. This limitation likely contributes to the inconsistent identification of low-abundant cardiac cell types across studies. While major cell populations such as cardiomyocytes, fibroblasts, and endothelial cells are consistently detected and represented across studies, rarer populations such as neurons, smooth muscle cells, pericytes, immune cells, adipocytes, and mesothelial epicardial cells can substantially vary (**Supplementary Figure 1E**). Misidentification of low-abundant cell types is likely caused by the limited number of nuclei recovered from these populations. It has been recently proposed that reliable identification of low-represented cell types during clustering requires a minimum of ∼ 500 nuclei or cells to achieve sufficient statistical power to distinguish between true biological signals and technical artifacts.^15^ Similarly, detection of clusters of low-represented cell types is heavily hampered by reduced nuclear RNA quality.^15^ Altogether, these observations highlight the need for an improved and standardized nuclei isolation strategy capable of maximizing nuclei yield while preserving RNA integrity and minimizing ambient RNA contamination.

**Table 1:**
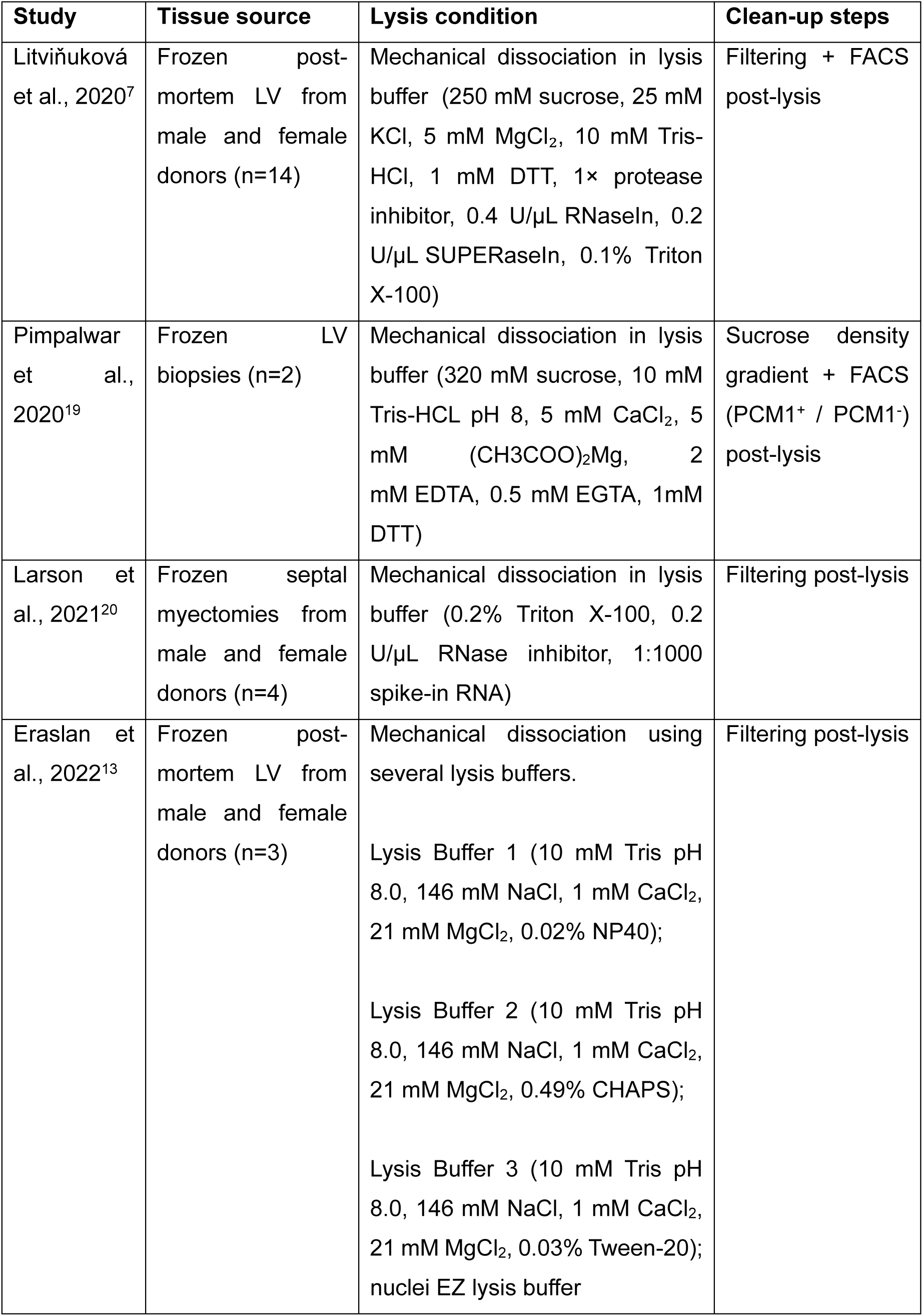
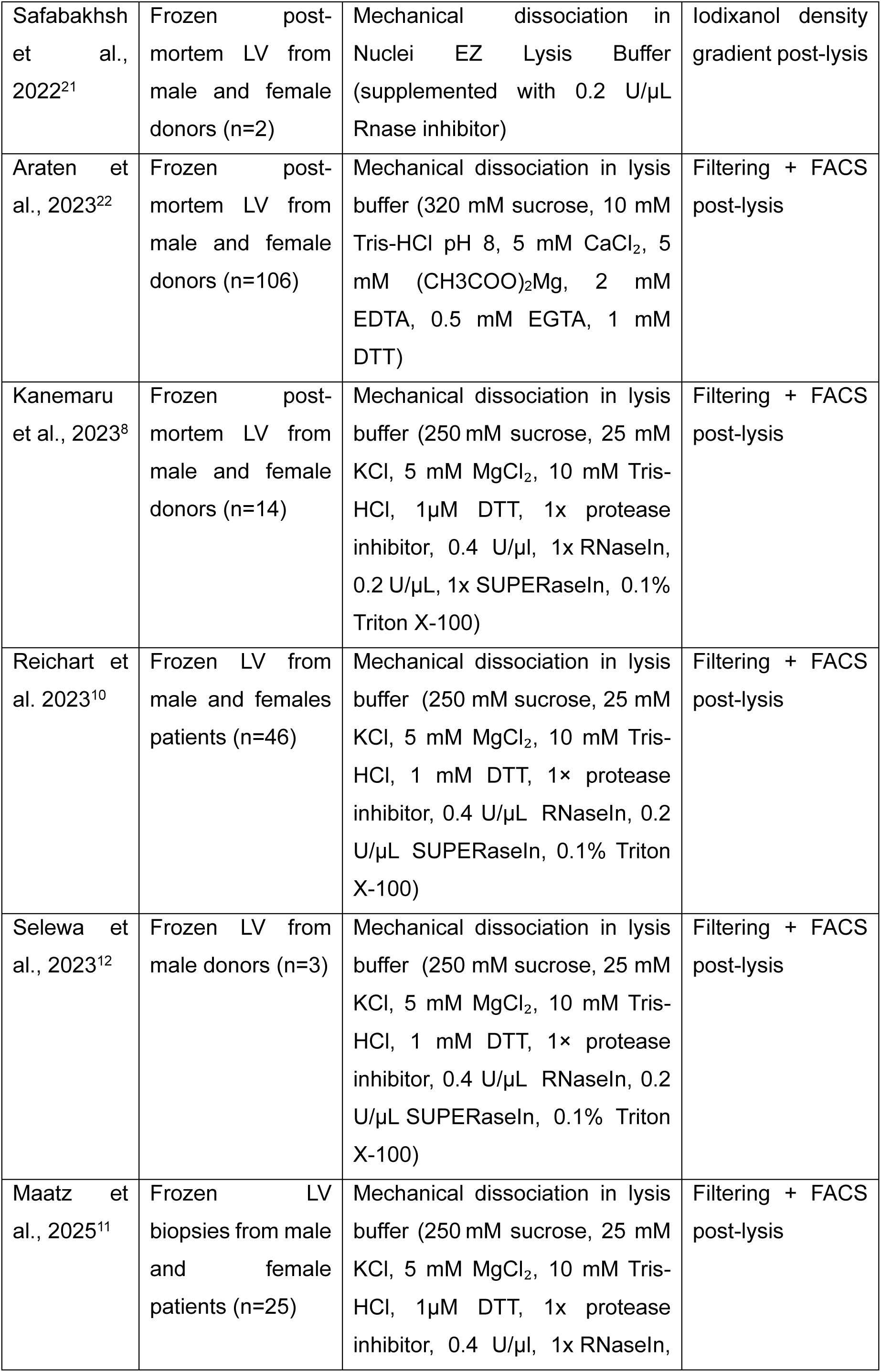
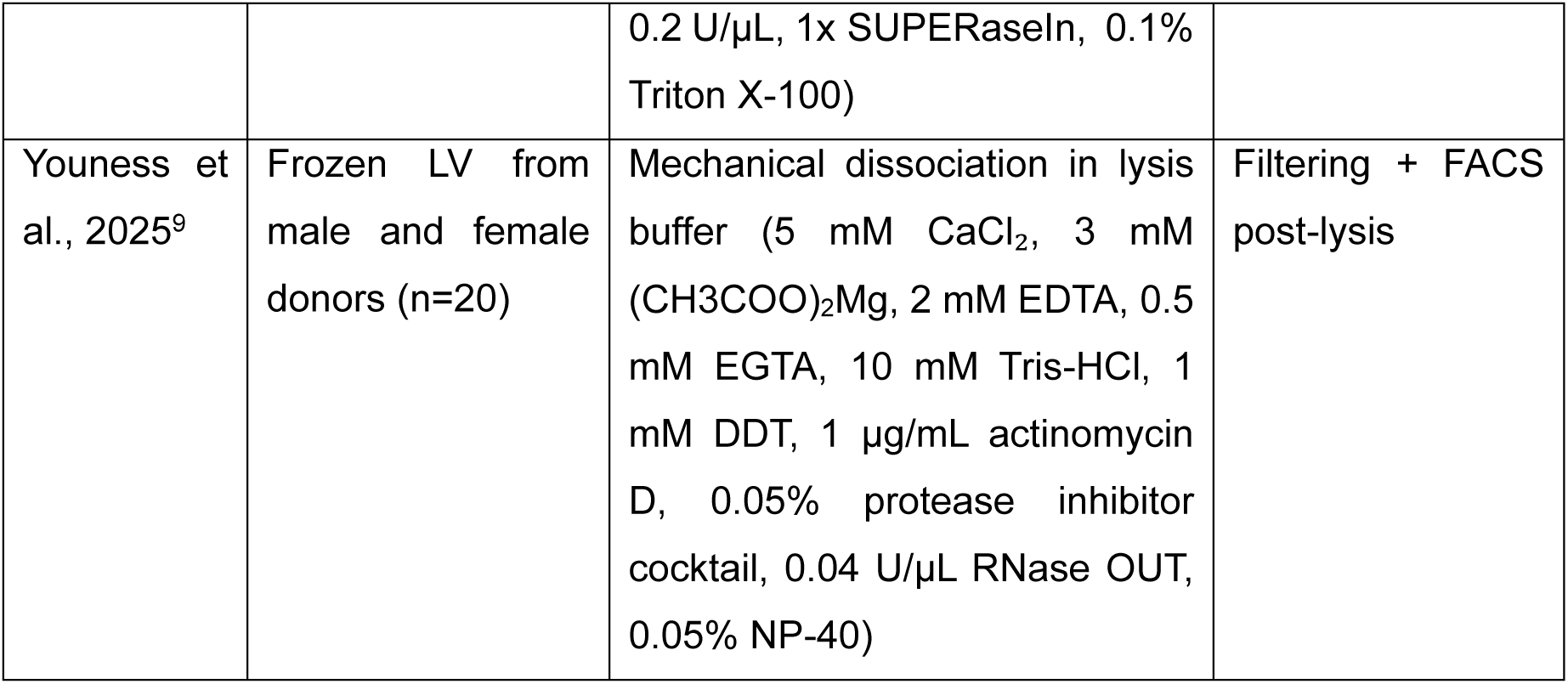
Nuclei isolation methods applied in previously published studies performing snRNA-seq on human adult left ventricle.

**Table 2:** Post-sequencing quality metrics of studies performing snRNA-seq from frozen adult human left ventricle.

| **Study** | **Nuclei loaded per reaction** | **Median n_Nuclei** | **Targeted sequencing depth per nucleus** | **Median n_Counts** | **Median n_Genes** | **Median pct_Counts_mt** |
| --- | --- | --- | --- | --- | --- | --- |
| Litviňuková et al., 2020<sup>7</sup> | 4000-10000 | 6937 | 20000-30000 reads | 2021 | 1080 | 1.42E-03 |
| Pimpalwar et al., 2020<sup>19</sup> | Curated dataset not publicly available |  |  |  |  |  |
| Larson et al., 2021<sup>20</sup> | Curated dataset not publicly available |  |  |  |  |  |
| Eraslan et al., 2022<sup>13</sup> | 7000 | 3444 | n/a | 923 | 781 | 0.00E+00 |
| Safabakhsh et al., 2022<sup>21</sup> | Curated dataset not publicly available |  |  |  |  |  |
| Araten et al., 2023<sup>22</sup> | Curated dataset not publicly available |  |  |  |  |  |
| Kanemaru et al., 2023<sup>8</sup> | n/a | 8566 | 20000-30000 reads | 2201 | 1164 | 1.18E-01 |
| Reichart et al. 2023<sup>10</sup> | n/a | 4415 | 30000-50000 reads | 2199 | 1296 | 8.96E-04 |
| Selewa et al., 2023<sup>12</sup> | 6000-8000 | 2412 | n/a | 3542 | 1635 | 5.54E-01 |
| Maatz et al., 2025<sup>11</sup> | n/a | 3409 | 30000-50000 reads | 2554 | 1404 | 5.47E-04 |
| Youness<br>et al.,<br>2025<sup>9</sup> | n/a | 7423 | 80000<br>reads | 7198 | 2723 | 0.00E+00 |

To address this need, we developed a hybrid nuclei isolation protocol for human cardiac tissue that integrates mechanical Dounce homogenization in Triton X-100-based lysis buffer, followed by sequential filtration, iodixanol density gradient, and FACS (**Figure 1**). The detergent-based mild lysis allows for efficient cellular disruption while preserving nuclear integrity, whereas density gradient purification removes big clumps of cellular debris that hinder FACS performance and prolong sorting time, conditions known to corroborate the degradation of the nuclear membrane. This integrated workflow consistently enables the isolation of high-quality nuclei suitable for robust and reproducible snRNA-seq from adult human heart tissue.

**Figure 1.**
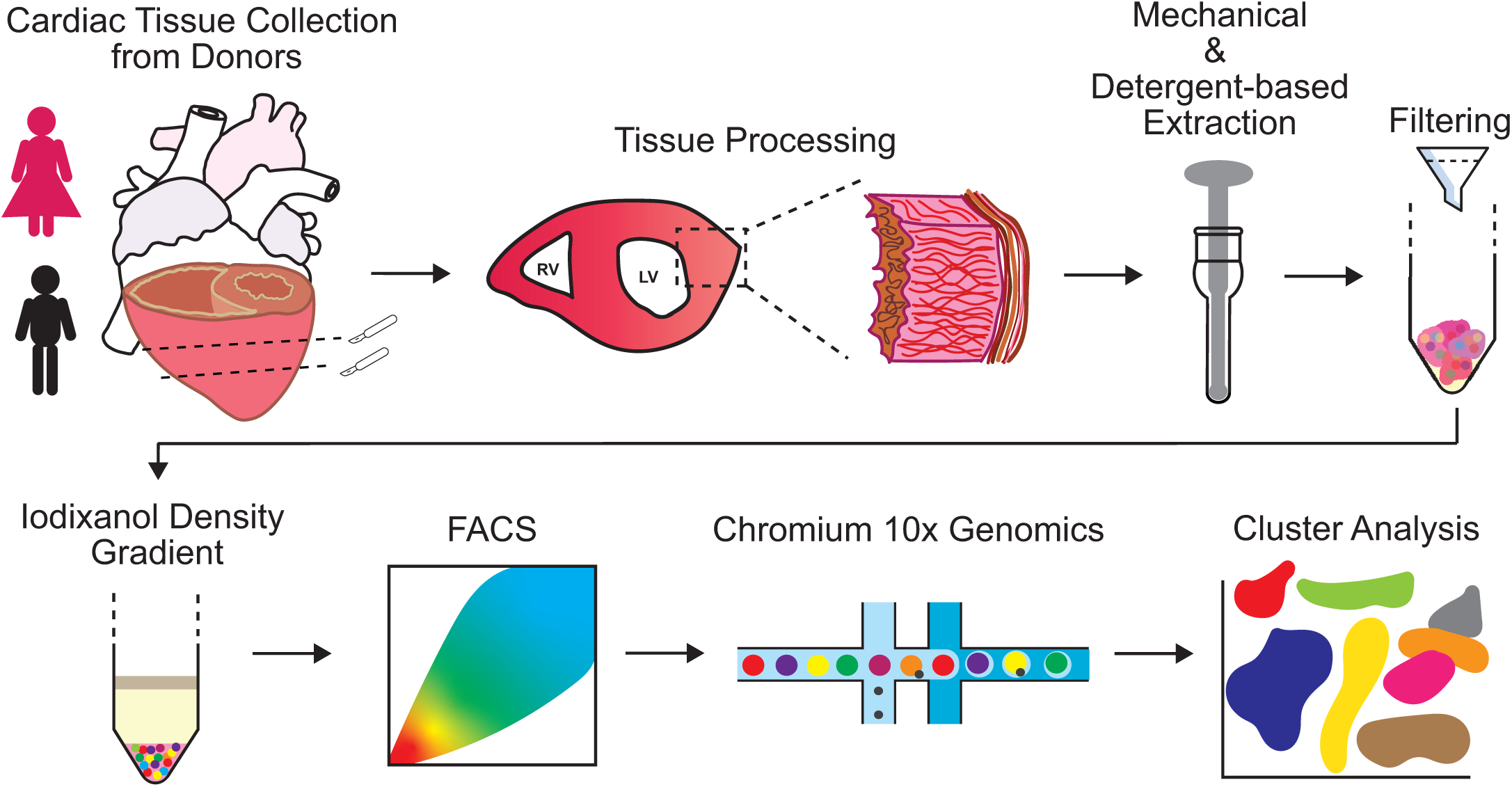
Hybrid nuclei isolation workflow optimized for human cardiac tissue. Human LV samples are collected and cryopreserved, then mechanically dissociated by Dounce homogenization in a detergent-based lysis buffer. The resulting lysate is filtered, and nuclei are purified using a sucrose density gradient followed by FACS clean-up for selection of DAPI^+^ nuclei. This protocol yields high-quality nuclei suitable for downstream snRNA-seq. RV=right ventricle; LV=left ventricle.

## MATERIALS

### LV tissue samples

**! CAUTION** Ethical approval from an institutional ethics committee may be required for handling and usage of human material. Personal protective equipment (PPE) should be used at all times during the experiment.

Approval for studies on human tissue samples was obtained from the Medical Ethics Committee of the University Medical Center Amsterdam, The Netherlands (METC-2024.0643). Donor hearts were collected with a post-mortem interval of 4-16 hours. Following thoracotomy, the aorta, the vena cava and the pulmonary veins were clamped, and the heart excised, washed in cold PBS, and systemically dissected on ice. The ventricles were isolated by a transverse cut performed below the atrioventricular valves. Two additional transverse cuts were then performed to generate three ∼ 2-cm-thick sections (ring A, ring B, and apex) (**Figure 2A-B**). Each transversal section was divided into nine transmural tissue blocks. Blocks corresponding to the section closest to the atrioventricular valves (ring A) were named blocks A1 to A9 (**Figure 2C**), whereas tissue blocks derived from the second transversal section, the closest to the apex (ring B), were named B1 to B9. Each tissue block was full-thickness sectioned into four portions, ensuring that each portion contained both the lumen trabeculae, the compact myocardium and the epicardium. Three portions were snap-frozen in liquid nitrogen and stored in −80 °C, and one portion was fixed in 4% PFA for 48 hours (**Figure 2D**). Unless otherwise stated, experiments were performed using transmural sections from block A2.

**Figure 2.**
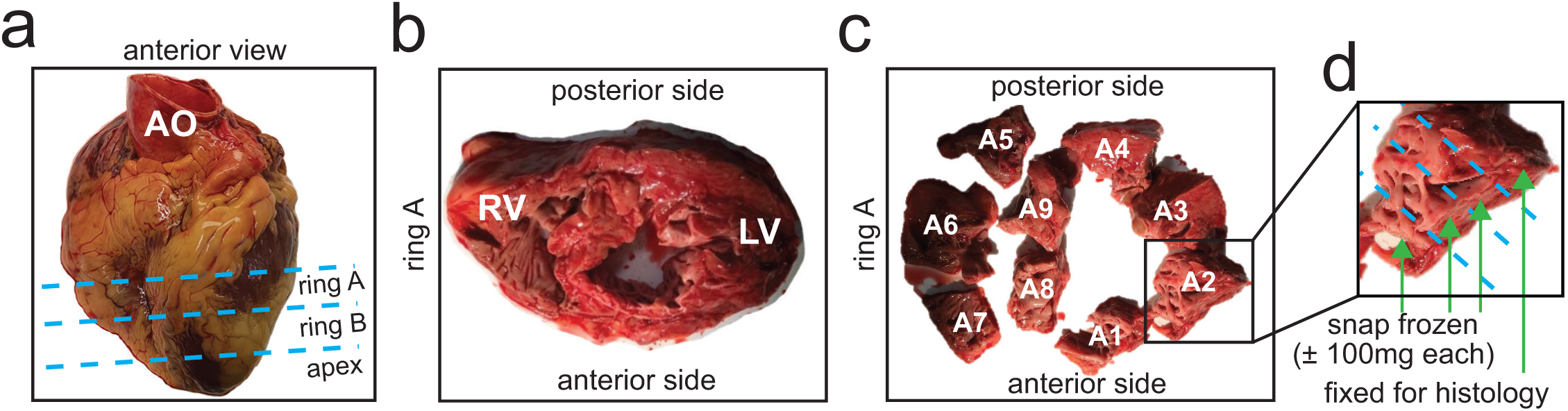
Workflow of human heart dissection. Following excision from the donor’s thorax, **(a)** the heart was oriented (anterior vs. posterior), and the ventricular chambers were isolated by cutting the heart below the atrioventricular valves. Two additional transverse cuts were performed to divide the ventricular chambers into three sections: Ring A, Ring B, and the apex. Dotted lines indicate the cutting planes. **(b)** For each section, anterior and posterior orientation was preserved, and the LV and RV were identified. **(c)** Ring A and Ring B were further subdivided into nine transmural tissue blocks (named A1 till A9) by additional cuts. Tissue blocks from Ring A are shown as an example. **(d)** Each tissue block was sectioned transmurally into four parts, with three portions snap-frozen in liquid nitrogen and one portion fixed in PFA for immunohistochemistry. AO=aorta; RV=right ventricle; LV=left ventricle.

**! CAUTION** Prolonged post-mortem interval prior to collection and snap freezing of the tissue can substantially compromise RNA quality. It is not recommended to use samples collected more than 6 hours post-mortem, as the RNA integrity number equivalent (RIN^e^) decreases markedly after this time frame (**Figure 3A-B**).

**Figure 3.**
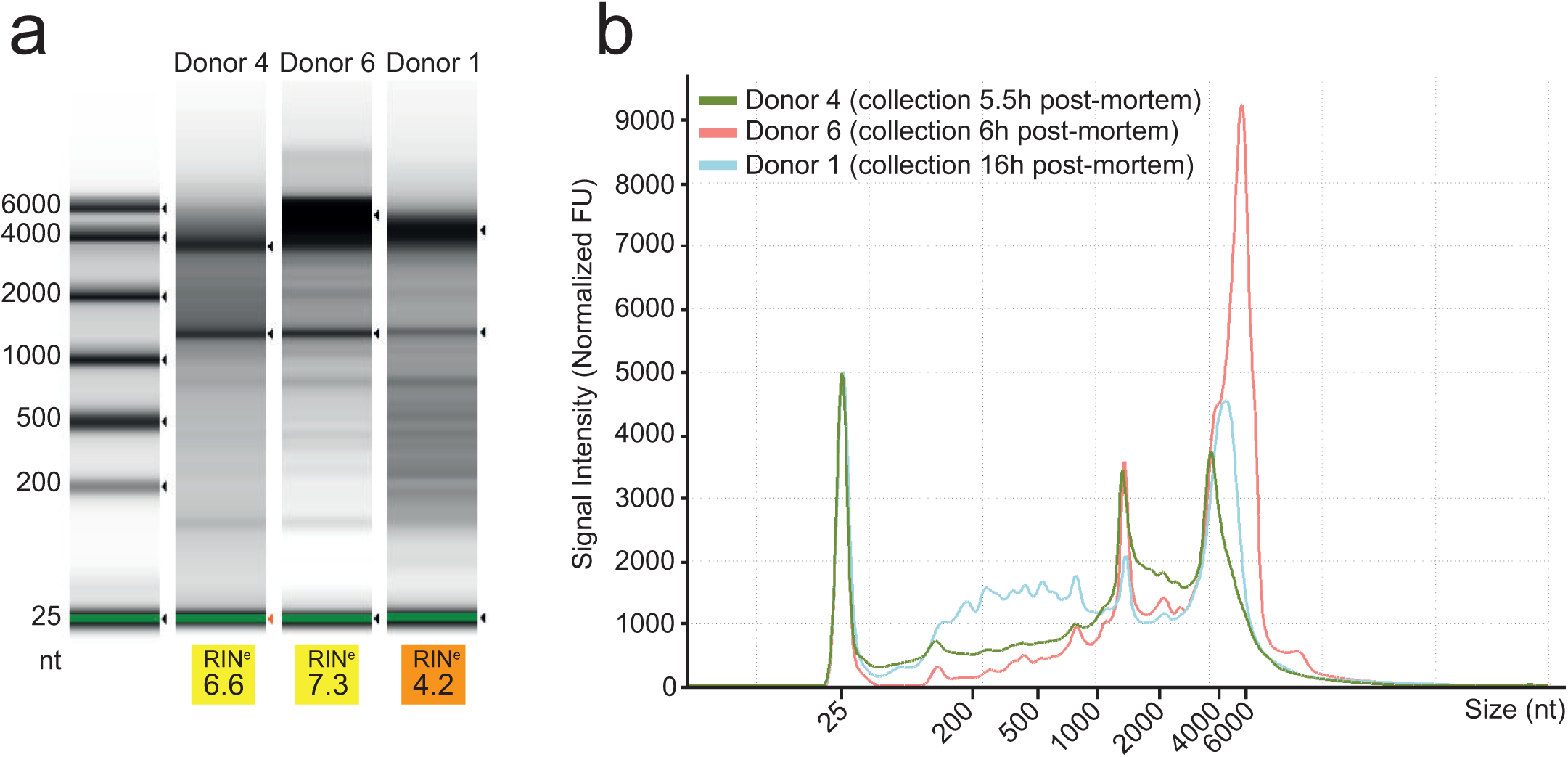
Relationship between post-mortem interval and RNA quality in human LV samples. **(a)** Representative TapeStation gel image with automated RNA quality assessment expressed as RIN^e^ score. Ribosomal RNA subunits are automatically detected by the software (black triangles next to each lane) and used to calculate the RIN^e^ score. Sample with a prolonged post-mortem interval (16 hours, donor 1) shows reduced RNA quality, as evidenced by multiple bands below the 18S rRNA band (∼1.5 kb), indicative of RNA degradation. **(b)** XY plot of signal intensity profiles derived from the TapeStation gel image. The *x-axis* indicates RNA fragment size (nt), and the *y-axis* indicates signal intensity (normalized fluorescent units). The peaks at ∼4.0 kb and ∼1.5 kb correspond to ribosomal RNA subunits 28S rRNA and 18S rRNA, respectively. The peak at ∼25 nt represents the TapeStation internal control. RIN^e^= RNA Integrity Number equivalent.

Demographic characteristics, post-mortem time of collection, and relevant clinical information of the donors included in this study are summarized in **Table 3**. Histological characterization of the LV samples is provided in **Supplementary Figure 2A**. Sirius Red (SR) staining demonstrates variable degrees of interstitial fibrosis (donor 4) or perivascular fibrosis (donor 5) in the samples used for snRNA-seq (**Supplementary Figure 2B**).

**Table 3:** Demographic characteristic, post-mortem time of collection and relevant cardiovascular and cancer clinical information of the donors.

| **Donor** | **Sex** | **Age**<br>**(years)** | **Post-**<br>**Mortem**<br>**Time**<br>**(hh:mm)** | **Cause of**<br>**Death** | **Smoker** | **Drinker** | **Relevant CV**<br>**Conditions and**<br>**Cancers** |
| --- | --- | --- | --- | --- | --- | --- | --- |
| Donor<br>1 | M | 78 | 16.00 | n/a | n/a | n/a | Anderoseptal infarct (2007),<br>cholangiocarcinoma (2012), gall bladder cancer (2012), small intestine cancer (2012), infarct LV (2020), cardiac asthma(2021), acute decompensation cardis with hypertension (2021) |
| Donor<br>2 | F | 76 | 4.30 | euthanasia | yes | yes | chronic stable angina, irregular blood pressure, implanted shunt in truncus coelicus , COPD phase 4 |
| Donor<br>3 | M | 97 | 4.00 | cardiac arrest | n/a | n/a | prostate cancer (2004), myocardial infarction (2016) |
| Donor<br>4 | M | 70 | 5.30 | respiratory depression | yes | n/a | Malignant broncus cancer with no compromised lung function (2021) |
| Donor<br>5 | M | 66 | 5.45 | n/a | n/a | n/a | n/a |
| Donor<br>6 | M | 85 | 6.00 | euthanasia | n/a | n/a | irregular heart beat<br>(2022), premature<br>ventricular complex<br>(2006),Tubular<br>Villous Adenoma<br>(1995 and 2009) |

### Reagents

4′,6-diamidino-2-phenylindole (DAPI, ThermoFisher, #D1306)

70% Ethanol

Bovine Serum Albumin (BSA, Sigma, #A8022)

Calcium Chloride dehydrate (CaCl_2_ • 2H_2_O, Merck, #1023821000)

DL-Dithiothreitol (DTT, Sigma, #D0632)

Dulbecco’s Phosphate-Buffered Saline (DPBS, ThermoFisher, #14190094)

Hydrogen chloride (HCl, Merck, #1003171000)

Magnesium Acetate tetrahydrate ((CH_3_COO)_2_Mg • 4H_2_O, Sigma, #M5661)

Magnesium Chloride hexahydrate (MgCl_2_ • 6H_2_O, Sigma, #44261)

Murine RNase inhibitor (NEB, #M0314)

Nuclease-free water (Qiagen Benelux, #129114)

OptiPrep^TM^ – 60% iodixanol (STEMCELL Technologies, #07820)

Potassium chloride (KCl, Sigma, #529552)

Potassium hydroxide (KOH, Merck, #1050331000)

RNase Zap solution (Invitrogen, #AM9780)

Sucrose (Sigma, #S0389)

Tricine (Sigma, #T0377)

Titriplex III^®^ - Ethylenediaminetetraacetic acid (EDTA, Merck, #1084180250)

Titriplex VI^®^ - Ethylene glycol-bis(2-aminoethylether)-N,N,N′,N′-tetraacetic acid (EGTA, Merck, #1084350025)

Triton X-100 (Sigma, #T8787)

Trypan Blue (Sigma, #T6146)

UltraPure Tris (ThermoFisher, #15504020)

## Equipment

2 mL Eppendorf tubes (Merck, #EP0030123344)

15 mL centrifuge tubes (Corning, #430791)

50 mL Falcon tubes (Corning, #352070)

7 mL Dounce tissue grinder with loose and tight pestles (Wheaton, #357542)

30 µm strainer (Sysmex, #04-004-2326)

100 µm strainer (Sysmex, #04-004-2328)

Agilent tapestation4200 - High Sensitivity D5000 (Agilent Technologies, #G2991AA)

Agilent tapestation4200 Reagents (Agilent Technologies, #RUO-5067-5593)

Cover glass for microscope slides (Meck, #BR470050)

Chromium Controller (10x Genomics, # PN-120270)

Chromium Next GEM Single Cell 3’ GEM, Library & Gel Bead Kit v3.1, 4 rxns (10x Genomics, PN-1000128)

Dry ice

DM6 B Upright Phase Contrast with 40X, 100X objective (Leica, #51701)

Glass beaker (Merck, # BR91236)

Ice

Styrofoam ice boxes

Magnetic stir bars

Magnetic stirrer

Microscope slides (VWR, #631-1164)

NovaSeq 6000 instrument (Illumina, #20012850)

NovaSeq S4 300-cycles Kit (Illumina #20046933)

Petri dish (Merck, #CLS430599)

pH meter

Refrigerated centrifuges with swing-out rotor (Hettich, #1554, #1706)

Round-bottom tube (Corning, #352001)

SH800S Cell Sorter (Sony, # SH800S)

Sterile surgical scalpel (ThermoFisher, #089275A)

Small tweezers

T10 Ultra-Turrax^®^ homogenizer (IKA, #0003737000)

Timer

Tubes shaker (Fisher Scientific, #NC2092426)

### BUFFERS PREPARATION ●TIMING 1 hour

**Preparation of Lysis (LYS) buffer**

For processing of four samples, prepare 125 mL of LYS buffer (20 mL per sample, plus excess for rinsing the Ultra-Turrax blades and the Dounce homogenizer). Scale-up volumes for processing of additional samples. To prepare 125 mL of LYS buffer:

1. Add the following to 121.25 mL nuclease-free water:

- 1.25 mL of 1 M Tris pH 8.0
- 0.625 mL 1 M CaCl₂
- 0.5 mL 0.5 M EDTA pH 8.0
- 0.625 mL 0.1 M EGTA pH 8.0
2. Mix the solution using a magnetic stirrer and autoclave.
3. Allow the solution to cool to room temperature, then add:

- 0.125 mL 1 M DTT
- 0.625 mL 1 M (CH3COO)_2_Mg.
4. Mix well and store at 4 °C for up to 4 days in the dark.

See also **Table 4** for the final buffer composition.

**Table 4:**
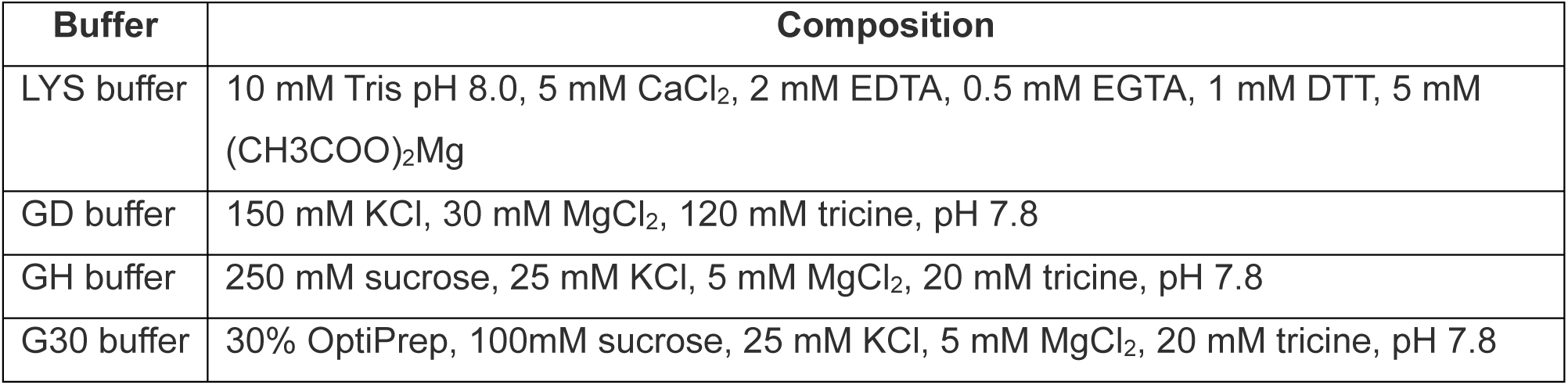
Composition of buffers required for hybrid nuclei isolation protocol.

**Preparation of 0.4% Triton-Lysis (T-LYS) buffer**

Prepare T-LYS buffer fresh on the day of the experiment.

For processing four samples, prepare 15 mL of T-LYS buffer:

1. Add 150 µL 40% Triton X-100 to 15 mL LYS buffer.
2. Mix on a magnetic stirrer until the detergent is fully dissolved.
3. Add 400U of murine RNase inhibitors per mL of T-LYS buffer.
4. Keep the buffer on ice until use.

**Preparation of G30 gradient (G30) buffer**

Prepare G30 buffer fresh on the day of the experiment.

For processing of four samples, prepare 12 mL of G30 buffer:

1. Mix the following components:

- 5 mL Optiprep (60% iodixanol)
- 1 mL of gradient diluent (GD) buffer (see composition below)
- 6 mL of gradient homologous (GH) buffer (see composition below)
2. Mix gently by inversion and keep on ice until use.

**Preparation of gradient diluent (GD) buffer**

1. Dissolve the following in 400 mL nuclease-free water:

- 5.59 g KCl
- 3.05 g MgCl_2_
- 10.75 g tricine
2. Adjust the pH to 7.8 using HCl and bring the final volume to 500 mL.
3. Filter-sterilize and store at room temperature for up to 2-3 months.

**Preparation of gradient homologous (GH) buffer**

1. Dissolve 10.27 g sucrose in 20 mL GD buffer.
2. Adjust pH to 7.8 and bring the volume to 120 mL with GD buffer.
3. Store at 4 °C for up to 1-2 months.

See **Table 4** for the final concentrations of all buffers listed in this section.

**Preparation of Staining (STAIN) buffer**

Prepare STAIN buffer fresh on the day of the experiment.

For processing four samples, prepare 5mL:

1. Dissolve 0.25g BSA in 5 mL of DPBS.
2. Mix until BSA is fully dissolved.
3. Add 100U of murine RNase inhibitors per 100 µL of STAIN buffer required.
4. Keep on ice until use.

**Preparation of DAPI Staining (DAPI-STAIN) buffer**

Prepare DAPI-STAIN buffer fresh on the day of the experiment.

For processing four samples, prepare 2mL:

1. Add 4 µL of DAPI solution (1 µg/µL) to 2 mL of STAIN buffer.
2. Mix gently and keep on ice until use.

### EQUIPMENT SETUP ●TIMING 30 minutes

Decontaminate all work surfaces with 70% EtOH and RNase decontaminant solution (RNase Zap). Fill a large ice box with ice. For each sample to be processed, precool a 7 mL Dounce tissue grinder, the pestles (placed in two 15 mL centrifuge tubes containing 5 mL LYS buffer), two 15 mL centrifuge tubes, one 50 mL Falcon tube, and two 2 mL Eppendorf tubes. Replace the caps of the two 15 mL centrifuge tubes with the 100 µm strainer and 30 µm strainer, respectively, and retain the original caps. Rinse the interior of the Dounce tissue grinder with 1-2 mL of LYS buffer to avoid friction during homogenization. Prepare a cleaning area for reusable equipment, including a beaker filled with nuclease-free water, a beaker filled with 70% ethanol, and a beaker with LYS buffer.

**▲ CRITICAL** Thorough cleaning of the 7 mL Dounce tissue grinder and the pestles and the Ultra-Turrax blades is fundamental to avoid sample cross-contamination, if they need to be reused for processing of multiple samples.

Prepare a second, smaller ice box. For each sample to be processed, precool a round-bottom tube containing 1 mL of T-LYS buffer. Prepare a tissue dissection area, placing a 15 cm Petri dish on ice and gathering a sterile surgical scalpel/razor blade, and small tweezers. Prepare a timer. Place the tube shaker in the cold room shaking at 45 rpm. Precool centrifuges with swing-out rotor at 4 °C. Collect the cardiac samples and keep them on dry ice until use.

### FACS GATING SETUP ●TIMING 1 hour

**▴ CRITICAL** The gating strategy should be optimized before the experiment.

FACS-based gating strategy is used to remove cell debris and purify nuclei. First, gate the instrument to select small particles based on forward- and side-scatter (**Figure 4A, left panel**). Next, gate the FACS instrument to sort only DAPI^+^ events (**Figure 4A, right panel**). Use an unstained control (no DAPI staining) to distinguish the negative population. DAPI^+^ events should then be filtered for the presence of aggregates, sorting only single nuclei (**Figure 4B**).

**Figure 4.**
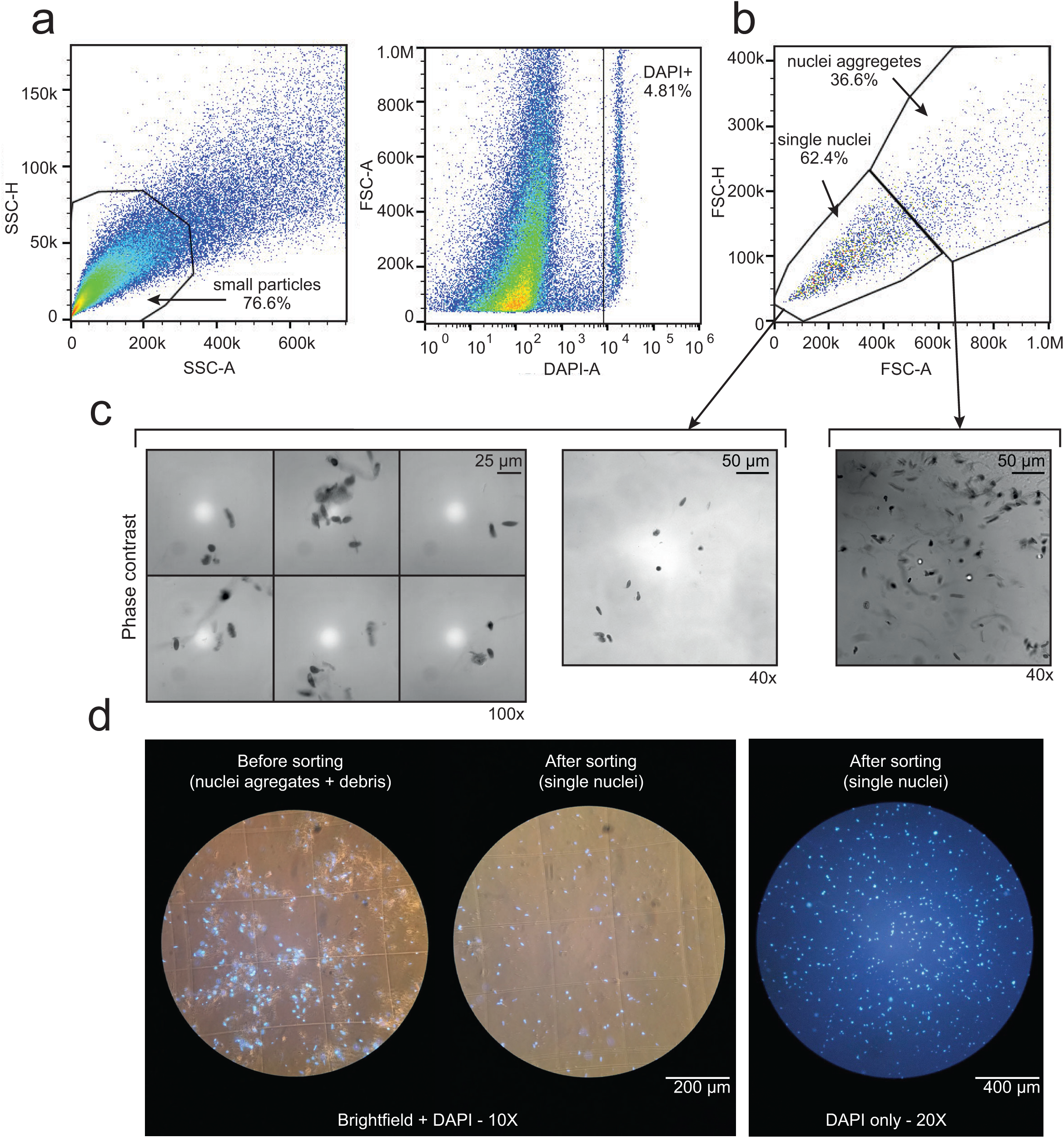
FACS gating setup and visual inspection of single nuclei sorted population. FACS instrument is first gated for **(a)** selection of small-sized particles (left panel), followed by selection of DAPI^+^ events (right panel). Ensure absence of nuclei aggregates by **(b)** gating to select only low-complexity events (single nuclei). **(c)** Gating strategy is validated by sorting both the single nuclei and nuclei aggregate populations and visually inspecting them by phase-contrast microscopy at high magnification. **(d)** On the day of the experiment, nuclei, debris distribution and presence of nuclei aggregates, before and after FACS clean-up can be quickly assessed by brightfield fluorescent microscope with low magnification. Scale bars are indicated in the figure. SSC=side scatter; FSC=Forward Scatter.

**▴ CRITICAL** Maintain the overall events rate below 2000-3000 events per second on the FACS instrument to minimize shear stress and detector saturation, which can compromise sorting accuracy. Incomplete tissue dissociation may cause clogging of the FACS instrument.TROUBLESHOOTING

During FACS gating setup, we recommend assessing nuclear membrane integrity and confirm proper gating strategy by sorting both the single-nuclei and aggregate populations, staining with Trypan Blue (1:1), and visually inspecting them using a phase-contrast microscope with a 40× and 100× objective (**Figure 4C**).

**▲ CRITICAL** Mount 5-10 µL Trypan Blue-stained nuclei solution on a microscope slide and cover with a coverslip. For phase-contrast microscopy, focus on the edge of the coverslip to identify the focal plane where nuclei are visible. Avoid extensive inspection of the samples using 40× or 100× phase-contrast prior to Chromium Controller loading, as locating the focal plane is time consuming, and prolong handling may compromise nuclear integrity. On the day of the experiment, a rapid assessment for debris and aggregates before and after FACS sorting can be performed using a brightfield microscope with 10× and 20× objective (**Figure 4D**).

### NUCLEI ISOLATION PROCEDURE

**Tissue processing and homogenization ●TIMING** 5 minutes per sample

1. Transfer the cardiac sample to the precooled Petri dish and dissect ∼0.10 g of tissue using a sterile surgical scalpel and fine tweezers (**Supplementary Figure 3A**). Make sure to dissect sample from lumen (trabeculae) to epicardium to avoid introducing bias in tissue’s cellular composition. Mince into smaller pieces and transfer to a precooled round-bottom tube containing 1 mL of T-LYS buffer. Start the timer.
2. Homogenize the tissue using the Ultra-Turrax for 60–120 seconds at max speed, until the suspension becomes homogenous (milkshake-like appearance).
3. Rinse the Ultra-Turrax blade with 1 mL T-LYS buffer and collect the rinse in the round-bottom tube.
4. Transfer the lysate to the 7 mL precooled Dounce tissue grinder, rinse the round-bottom tube with an additional 1 mL of T-LYS, and transfer it to the Dounce tissue grinder.
5. Homogenize with 10 strokes using the loose pestle, followed by 15 strokes with the tight pestle.
6. Transfer the lysate to a 50 mL tube and shake for 10 minutes at 4 °C on a tube shaker to facilitate full separation between the nuclei and the tissue fibers.

**▲ CRITICAL** Thoroughly clean the Ultra-Turrax blades, Dounce tissue grinder, and the pestles with nuclease-free water, 70% ethanol, and LYS buffer before processing a new sample to prevent cross-contamination.

7. Repeat steps 1-6 for each sample, maintaining identical processing times.

**Nuclei purification ●TIMING** 40 minutes

8. Add 3 mL precooled LYS buffer to the T-LYS buffer containing the nuclei to reduce the Triton X-100 concentration and filter the lysate through a 100 µm strainer. If clogging occurs, gently move the debris around using a pipette tip.
9. Wash the strainer with 2 mL precooled LYS buffer before discarding it.
10. Filter the lysate through a 30 µm strainer and wash with 2 mL LYS buffer. If clogging occurs, gently move the debris using a pipette tip.
11. Wash the strainer with 2 mL precooled LYS buffer before discarding it.
12. Centrifuge the lysate at 1000 × g for 5 minutes at 4 °C using a precooled centrifuge with swing-out rotor.
13. Carefully discard the supernatant, leaving the nuclei and small debris pellet intact (**Supplementary Figure 3B**).

**▲ CRITICAL** Prior to experiment, ensure that the tissue lysis does not compromise the nuclear membrane integrity. Pellet of nuclei and debris can be resuspended in STAIN buffer, mixed with Trypan Blue solution (1:1), and visually inspected using a phase-contrast microscope with a 40× and 100× objective (**Supplementary Figure 3C**). At this stage, presence of debris is normal. If nuclear membranes are compromised, consider reducing lysis time. **? TROUBLESHOOTING**

14. Resuspend the pellet in 300 µL of G30 buffer and transfer it to a 2 mL Eppendorf tube.
15. Carefully underlay 1 mL of G30 buffer, allowing the G30 buffer containing nuclei and small debris to form a clear layer above the clean G30 buffer.
16. Centrifuge the sample at 8000 x g for 20 minutes at 4°C using a precooled centrifuge with a swing-out rotor.
17. Completely discard the supernatant and gently wash the pellet with 500 µL STAIN buffer.TROUBLESHOOTING
18. Resuspend the pellet in 250 µL of DAPI-STAIN buffer.

**▲ CRITICAL** Incomplete removal of the G30 buffer might interfere with the droplet formation during FACS, requiring new instrument calibration. **? TROUBLESHOOTING**

**Sorting and counting of nuclei ●TIMING** 1 hour

19. Sort at least 100000 DAPI^+^ events into collection tubes containing 50 µL STAIN buffer.

**▲ CRITICAL** Sorting more events will increase sorting time, which can compromise quality of nuclei.

20. Transfer the sorted nuclei to a precooled 2 mL Eppendorf tube and centrifuge at 1000 × g for 10 minutes at 4 °C, using a precooled centrifuge with a swing-out rotor.
21. Carefully remove 40-70% of the supernatant and gently resuspend to obtain a final concentration of ∼ 1000–2000 nuclei/µL.

**▲ CRITICAL** A visible pellet is typically obtained when sorting more than 250000 DAPI^+^ events. For lower yields, volume reduction has to be performed without visual guidance as no pellet will be visible. Additionally, the precise percentage of volume to be discarded to reach the desired concentration depends on the volume increase produced by the FACS sorter and the accuracy of sorting. This must be determined prior to the experiment.

22. Mix 10 µL of nuclei suspension with Trypan Blue solution (1:1) and determine nuclei concentration using an automated cell counter.

**▲ CRITICAL** Automated cell counter may underestimate nuclei concentration due to focal-plane errors. Manually inspect the focal plane used for the counting. **? TROUBLESHOOTING**

### POST- ISOLATION WORKFLOW ●TIMING 3-7 days

In our experimental setup, we isolated nuclei using either our hybrid method or a commonly applied two-step clean-up strategy (**Figure 5**), similar to that used in most studies listed in **Table 2**. Four adult human LV samples were processed using the hybrid protocol, and one human adult LV sample was processed using the two-step method (**Figure 5**). We next compared post-sequencing metrics from snRNA-seq performed on nuclei isolated by these two approaches to evaluate the robustness of our hybrid protocol.

**Figure 5.**
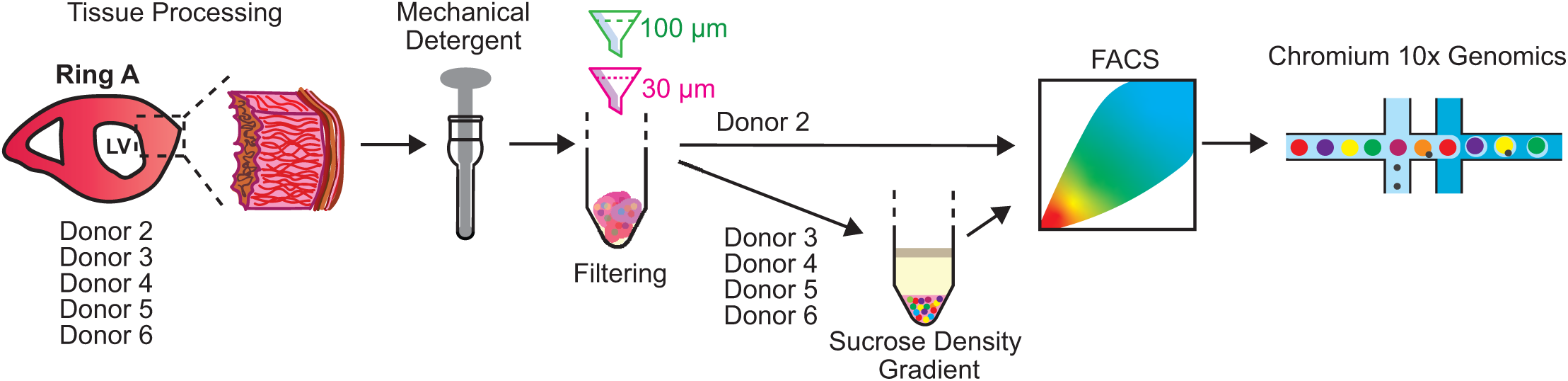
Experimental setup. Schematic overview of the workflow followed. After tissue dissociation and filtering, human LV samples were processed using the two-step clean-up approach employed for nuclei isolation (filtering and FACS) or were processed according to our hybrid protocol (filtering, sucrose density gradient, FACS) prior to snRNA-seq. LV=left ventricle; FACS= fluorescence-activated cell sorting.

#### 10x library preparation and sequencing

For each preparation, a volume containing 33000-44000 nuclei per reaction was loaded into a Chromium Next GEM Single Cell 3’ GEM Chip. 3′ gene expression libraries were generated according to the manufacturer’s specifications of the v3.1 Chromium Next GEM Single Cell Reagent Kit.

**▪ PAUSE POINT** Quality controls on cDNA can be performed later. cDNA can be stored at - 20 °C until further processing.

cDNA quality was assessed using TapeStation High-Sensitive DNA analysis. Libraries from four reactions were pooled together and sequenced using Illumina NovaSeq 6000 (S4 300-cycles kit), targeting to recover 80 million paired-end reads per sample.

#### Mapping

Cell Ranger 7.0.1. was used for read processing and alignment. Raw sequencing data (FASTQ files) were mapped to the human reference genome 2020-A (GRCh38, GENCODE v32/Ensembl 98), provided by 10x Genomics, using the CellRanger v6.0.2 pipeline, with intronic reads included (--include-introns=TRUE) in the count matrix.

**▲ CRITICAL** In Cell Ranger pipeline prior to v7.0, the --include-introns option is disabled by default. Enabling introns inclusion increases transcriptome mapping rate by ∼ 40% (**Supplementary Figure 4**), and is essential for snRNA-seq, as a substantial fraction (∼ 50%) of nuclear RNAs include intronic sequences.^16^

#### Post-sequencing quality metrics prior to data filtering

Analysis of n_Nuclei demonstrated that the hybrid workflow consistently outperformed the two-step method, yielding approximately fourfold higher nuclei recovery (**Table 5**; **Supplementary Figure 5A**). The distribution of n_Counts indicated that the hybrid protocol approximately doubled the number of detected UMIs per nucleus (**Table 5**, **Supplementary Figure 5B**), suggestive of substantially enhanced transcript capture. Importantly, this increase was not associated with elevated mitochondrial RNA content, as pct_Counts_mt remained comparable between the two approaches (**Table 5**, **Supplementary Figure 5C**). Consistently, gene detection was enhanced, with an average of ∼ 1500 genes detected per nucleus using the hybrid protocol (**Table 5**, **Supplementary Figure 5D**). Notably, this level of gene coverage was achieved at a sequencing depth of ∼ 8000-10000 reads per nucleus, whereas most studies listed in **Table 2** required substantially higher sequencing depth (two to five times higher) to reach similar number of genes.

**Table 5.**
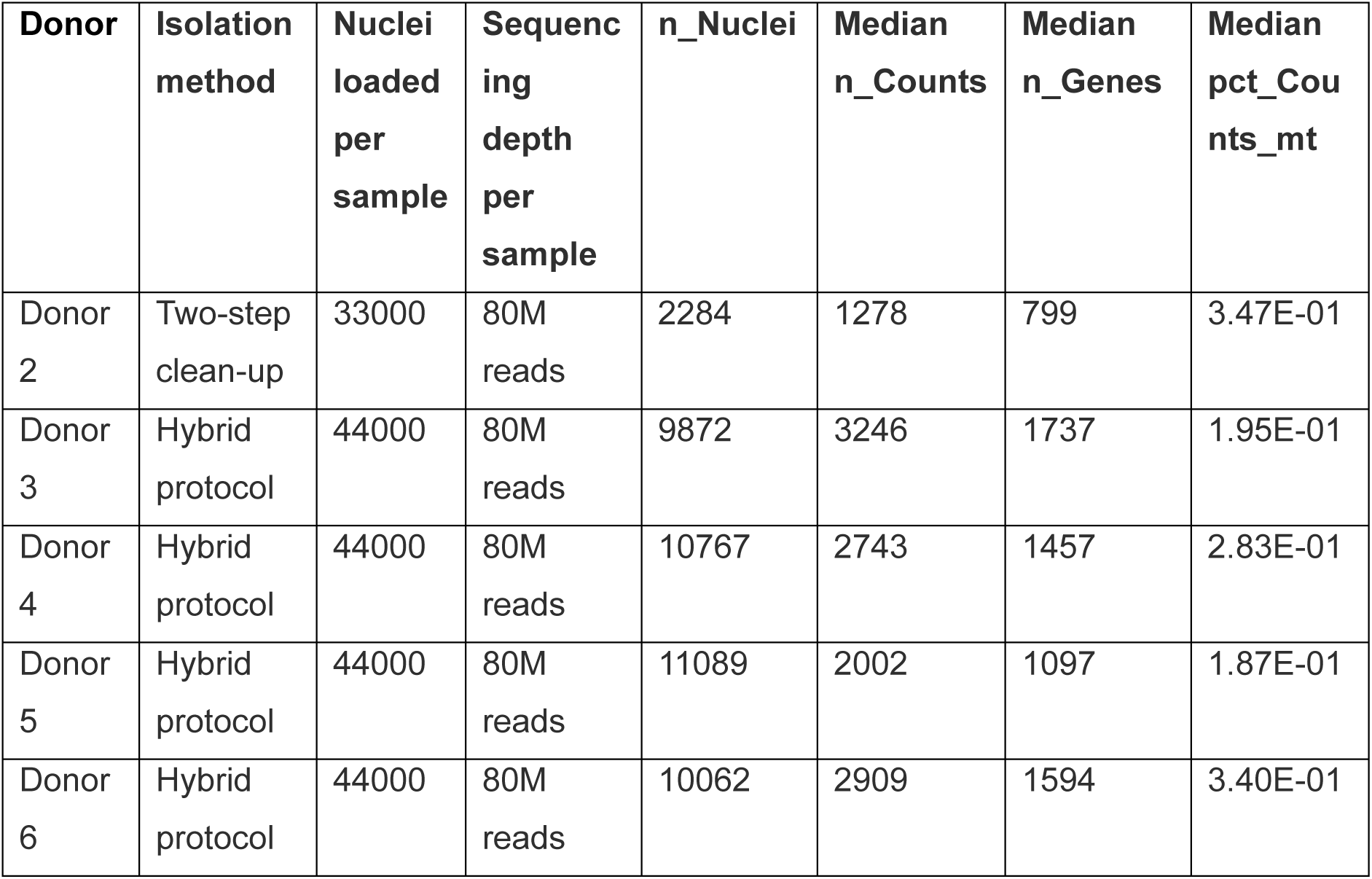
Post-sequencing quality metrics for snRNA-seq from nuclei isolated from frozen adult human left ventricle using either the two-step clean-up approach or the hybrid isolation protocol.

| **Donor** | **Isolation method** | **Nuclei loaded per sample** | **Sequencing depth per sample** | **n_Nuclei** | **Median n_Counts** | **Median n_Genes** | **Median pct_Counts_mt** |
| --- | --- | --- | --- | --- | --- | --- | --- |
| Donor 2 | Two-step clean-up | 33000 | 80M reads | 2284 | 1278 | 799 | 3.47E-01 |
| Donor 3 | Hybrid protocol | 44000 | 80M reads | 9872 | 3246 | 1737 | 1.95E-01 |
| Donor 4 | Hybrid protocol | 44000 | 80M reads | 10767 | 2743 | 1457 | 2.83E-01 |
| Donor 5 | Hybrid protocol | 44000 | 80M reads | 11089 | 2002 | 1097 | 1.87E-01 |
| Donor 6 | Hybrid protocol | 44000 | 80M reads | 10062 | 2909 | 1594 | 3.40E-01 |

**Table 6:** Troubleshooting.

| Step | Problem | Possible reason | Solution |
| --- | --- | --- | --- |
| FACS instrument setup | Populations shifting during FACS sorting | Clogging of FACS instrument due to high number of nuclei aggregates<br>High events rate | Extend time of Ultra-Turrax and Dounce homogenization<br>Reduce sample pressure |
| Step 13 | No nuclei are observed using phase-contrast | Nuclei suspension is overdiluted | Pellet the nuclei and debris suspension by centrifugation and resuspend in lower volume |
| Step 17 | No pellet is visible at the bottom of the Eppendorf tube | Low number of nuclei due to low starting material<br>Incomplete lysis<br>Wrong layering of G30 buffer (step 15)<br>Use of centrifuge with a fixed-angle rotor (pellet on the side of the tube) | Increase amount of starting material<br>Increase lysis time<br>Repeat G30 buffer layering using pipette tips with lower bore diameter<br>Adopt a centrifuge with a swing-out rotor |
| Step 18 | Loss of calibration of FACS instrument | Residual presence of G30 buffer | Consider performing additional washes at step 17 to avoid G30 buffer contamination |
| Step 22 | Low number of nuclei detected | Wrong focal plane of the cell counter<br>Cell counter is set to exclude low-sized particles | Manually adjust the focal plane till nuclei are in focus and repeat quantification<br>Edit cell counter parameters allowing inclusion of particles with minimal size of 3-5 $\mu\text{m}$ |

#### Data filtering

Sequencing data from all samples were analyzed in R (version 4.2.2) and processed using the Seurat package (version 4.3.0.1 – *Hao et al., 2021* ^17^). Nuclei with more than 3% mitochondrial reads were excluded. Mitochondrial genes were removed from the expression matrix to eliminate transcripts originating from outside the nucleus. Nuclei with low genes or UMI counts were removed. **F**iltering led to the removal of 14% of nuclei input derived from Donor 2, while only 2-5% of the nuclei input derived from all the other donor was removed. The dataset was normalized for sequencing depth per nucleus and log-transformed using a scaling factor of 10000. Principal component analysis (PCA) was performed on the 2000 most variable features, and the first 29 principal components (PCs) were used to generate the UMAP and find clusters. Batch effects between libraries were corrected using the Harmony package (version 0.1.1). For the identification of potential doublets, an initial clustering was performed using Louvain clustering based on a shared nearest neighbour graph, as implemented in the FindClusters function in Seurat with parameters k.param = 5 and resolution = 0.3. These clusters were used to identify potential doublets (doublets.score ∼ 0.999) using the scDblFinder,^18^ package v1.12.0 (**Supplementary Figure 5E**). Across all samples, we identified less than 10% potential doublets, indicating that most droplets contained single nuclei (**Supplementary Figure 5F**). This percentage of doublets is consistent with what to be expected for the number of nuclei recovered, as specified in the Chromium Next GEM Single Cell 3’ v3.1 user guide (CG000204 Rev D). Nuclei labeled as doublets were excluded from downstream analysis using scDblFinder.

#### Clustering and differential expression

Clustering was performed on the filtered dataset with parameters k.param = 10, resolution = 0.3. Differentially expressed genes (DEGs) were identified for each cluster using the Wilcoxon rank-sum test implemented in the FindAllMarkers function in Seurat, with parameters logfc.threshold = 0.25, return.threshold = 0.01, only.pos = TRUE, min.pct = 0.25. This analysis identified 17 distinguished clusters corresponding to 11 cardiac cell types, which were annotated based on established marker gene expression (**Figure 6A**). The top 10 DEGs per cluster are shown in **Supplementary Figure 6**. Examination of the contribution of each individual sample to the 17 identified clusters showed that after nuclei isolation using the two-step clean-up approach (donor 2), nuclei were predominantly derived from fibroblasts (cluster 0), cardiomyocytes (clusters 1 and 11), myeloid cells (cluster 6), and adipocytes (cluster 14). In contrast, samples processed using the hybrid isolation protocol contributed to broader recovery of different cardiac cell populations (**Figure 6B**). Consistently, the two-step protocol yielded a biased cellular composition with the overrepresentation of cardiomyocytes, compared to what yielded by the use of the hybrid protocol (**Figure 6C**).

**Figure 6.**
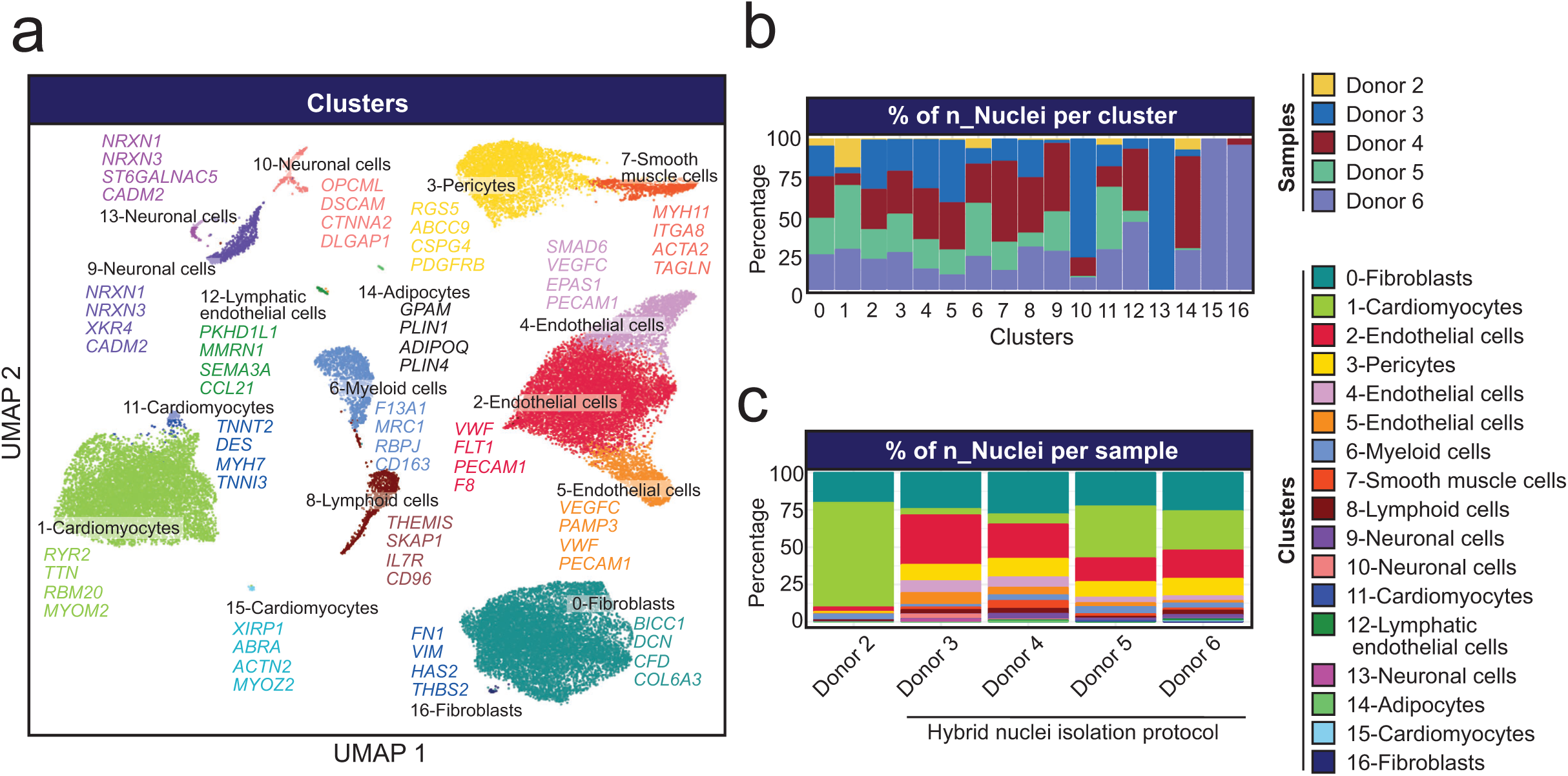
Cluster analysis and cell type identification in human adult LV samples. **(a)** UMAPs embeddings of the of pooled data from all sequenced samples (41356 nuclei), highlighting identification of multiple cardiac cell types. Cell specific markers used for cluster annotation are depicted in the figure. **(b)** Bar chart showing, the contribution (% of nuclei) of each sample to each cluster. **(c)** Bar chart showing the cluster distribution per sample. Samples processed using hybrid protocol are indicated. UMAPs=uniform manifold approximation and projection

## CONCLUSIONS

Single-nucleus RNA-seq has become an essential approach for investigating cardiac biology and disease and is now routinely used in both fundamental and translational studies. However, the quality and integrity of isolated nuclei remain a major bottleneck of this technique, often resulting in low gene detection, biased cell-type representation, and substantial inter-study variability. To address these limitations, we describe a hybrid nuclei isolation that integrates multiple sequential clean-up steps to improve nuclei recovery and preserve RNA integrity from frozen adult human heart tissue. This approach consistently yields higher nuclei recovery and improved transcript capture without increasing mitochondrial RNA contamination. Adoption of this hybrid protocol may help to significantly reduce the heterogeneity commonly observed across snRNA-seq studies.

## TIMING

Buffers preparation: ∼1 hour

Equipment setup: ∼30 minutes

FACS gating setup: ∼1 hour

Tissue processing and homogenization (Steps 1-7): ∼5 minutes per sample

Nuclei purification (Steps 8–18): ∼40 minutes

Sorting and counting of nuclei (Steps 19-22): ∼1 hour (including 10 minutes per samples for FACS sorting)

Post-isolation workflow: ∼3-7 days

## ACKNOWLEDGEMENTS

This work is supported by Dr. Dekker Senior Scientist Fellowship from the Dutch Heart Foundation (NHS2020T041 to M.M.G) and the Out Of The Box Grant from Amsterdam Cardiovascular Sciences.

## AUTHORS’ CONTRIBUTION

R.C. and M.M.G. designed the experiments. R.C. and R.B. performed all experiments. R.C., R.B., R.A.B. and M.M.G. analyzed the data. J.H., R.J.O., M.J.B.vd.H provided materials. R.C., R.B., R.A.B. and M.M.G. wrote the manuscript. All authors revised the manuscript.

## COMPETING FINANTIAL INTERESTS

The authors declare no competing financial interests

---

**Supplementary Figure 1.**
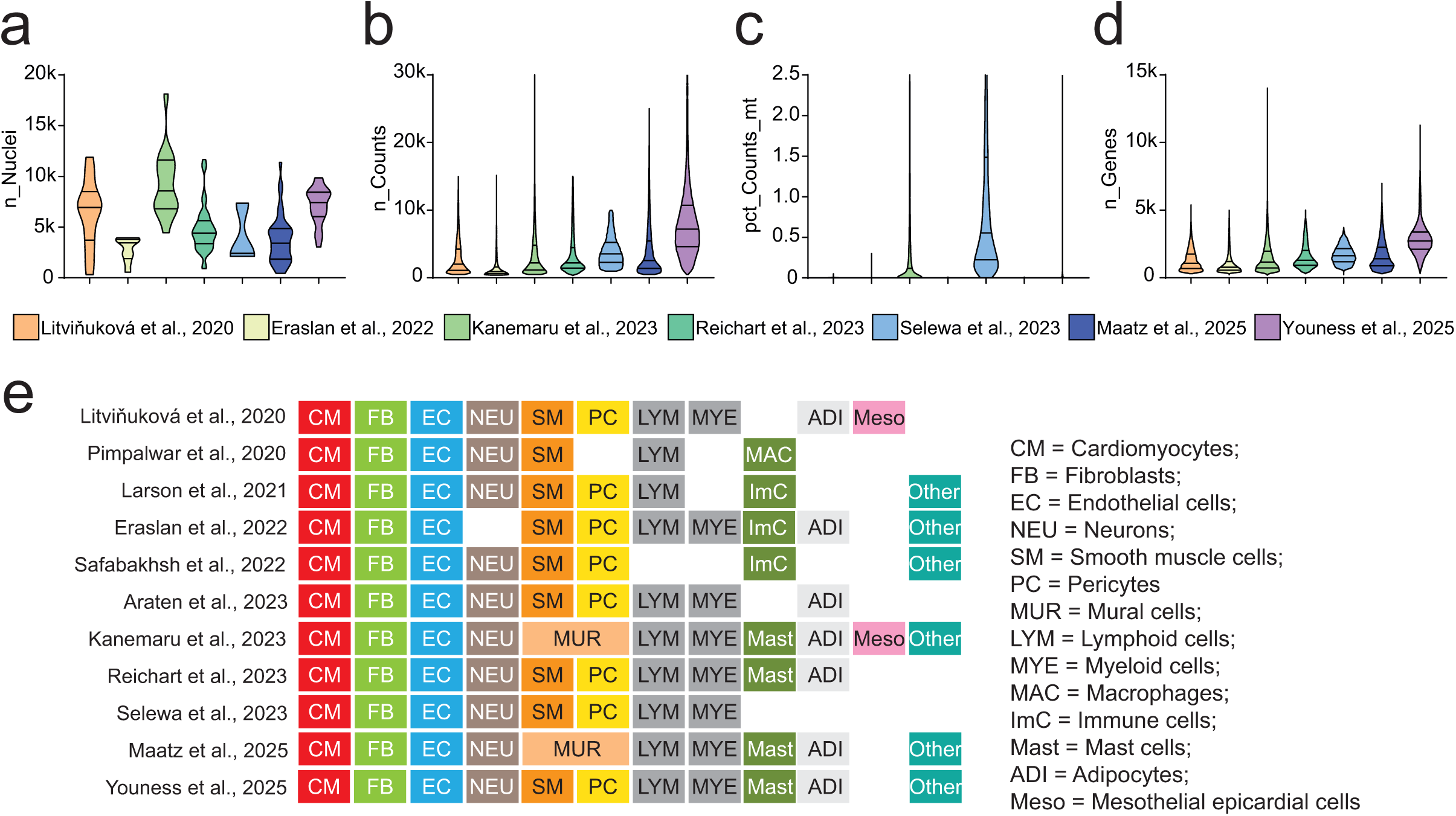
Post-sequencing quality metrics and cell type annotation across snRNA-seq studies of human LV. **(a–d)** Violin plots summarizing post-sequencing quality metrics across studies: **(a)** number of nuclei recovered per study (n_Nuclei), **(b)** median of UMIs per nucleus (n_Counts), **(c)** median percentage of reads mapping to mitochondrial RNA (pct_Counts_mt), and **(d)** median of genes detected per nucleus (n_Genes). For each distribution, the median and interquartile range are indicated by solid lines. Analyses include only studies that publicly released curated data. **(e)** Cell type annotations reported by individual studies performing snRNA-seq on human LV samples. The identification of low-abundance cell populations varies markedly across studies.

**Supplementary Figure 2.**
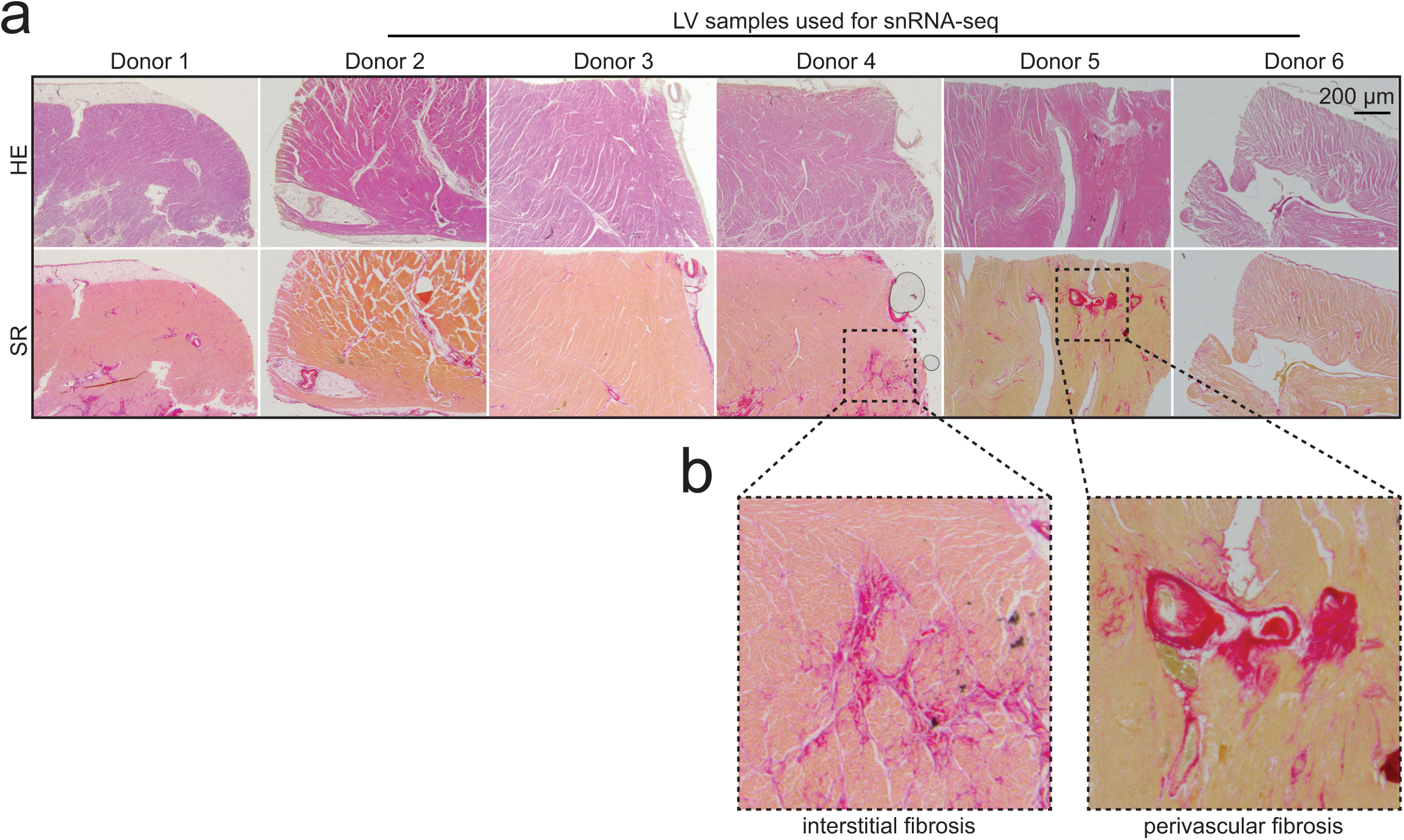
Histological characterization of the LV samples used for this study. **(a)** Representative images of LV sections stained with Hematoxylin and Eosin (H&E) and Sirius Red (SR). Samples selected for snRNA-seq (indicated in the figure) display varying degrees of cardiac fibrosis. **(b)** Higher-magnification of SR-stained sections highlighting interstitial and perivascular fibrosis observed in two of the five samples used for snRNA-seq. Scale bar is indicated in the figure.

**Supplementary Figure 3.**
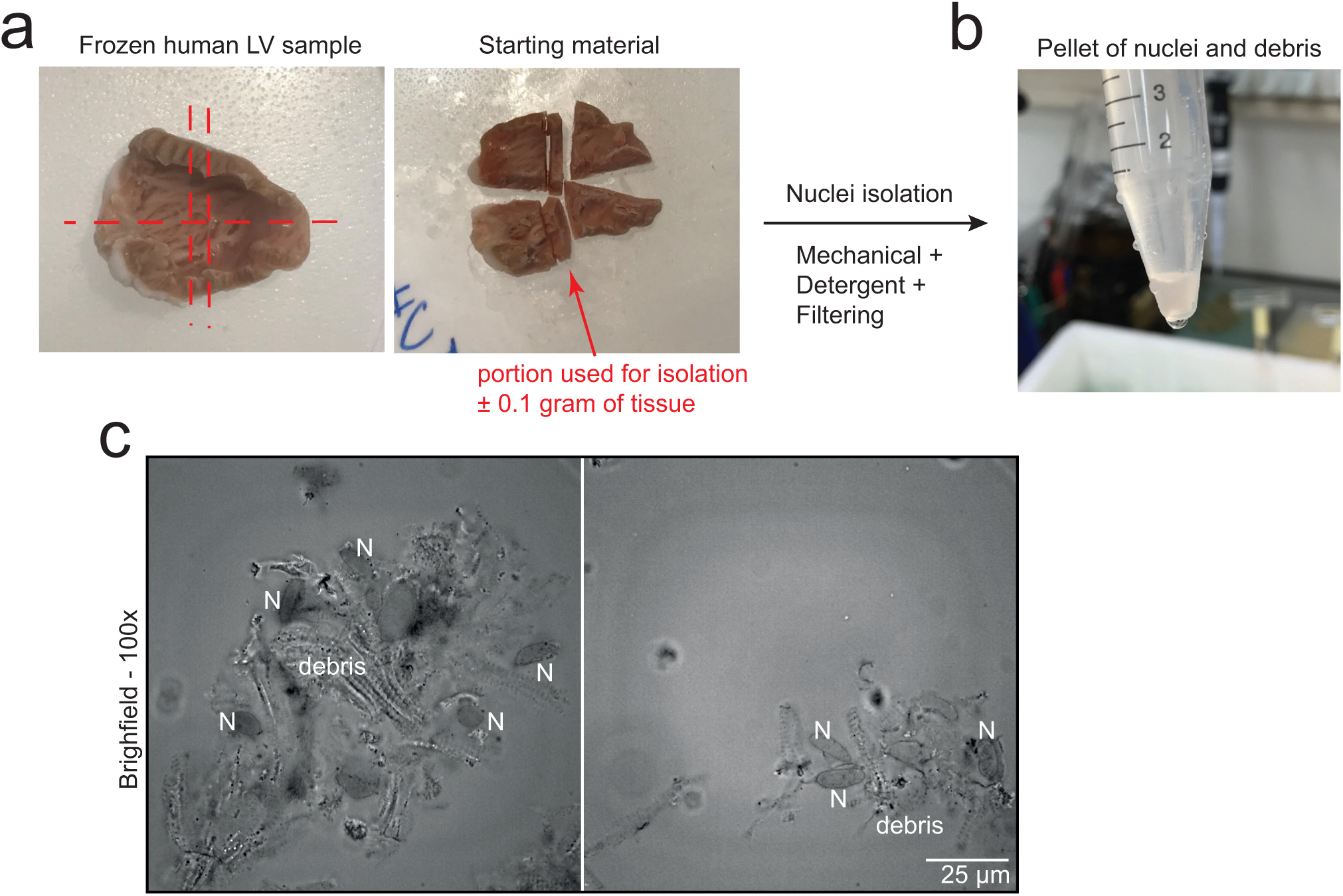
Tissue processing and post-lysis assessment of nuclei integrity. **(a)** 100 mg of cardiac sample can be isolated on ice and used for homogenization. Following filtration and centrifugation, **(b)** a pellet of nuclei and debris appears. Ensure that lysis conditions did not alter nuclear integrity by staining the sample of nuclei and debris with Trypan blue, inspecting **(c)** the sample using a phase contrast microscope with high magnification. Scale bar is indicated in the figure. LV=left ventricle; N=nuclei.

**Supplementary Figure 4.**
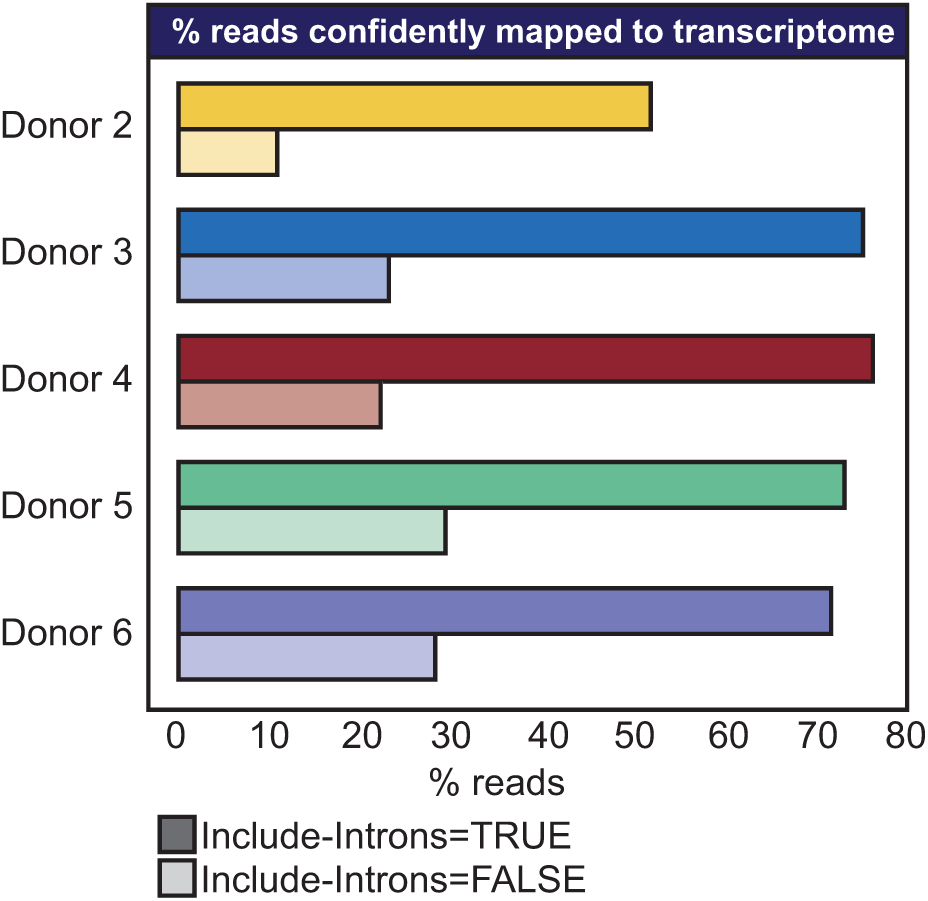
Effect of intronic read inclusion on snRNA-seq transcriptome mapping rate. Bar chart showing the percentage of confidently mapped reads to the transcriptome per sample when intronic reads are included (bars with solid colours) or excluded (bars with semi-transparent colours). Inclusion of intronic reads increases the overall mapping rate by approximately 40%.

**Supplementary Figure 5.**
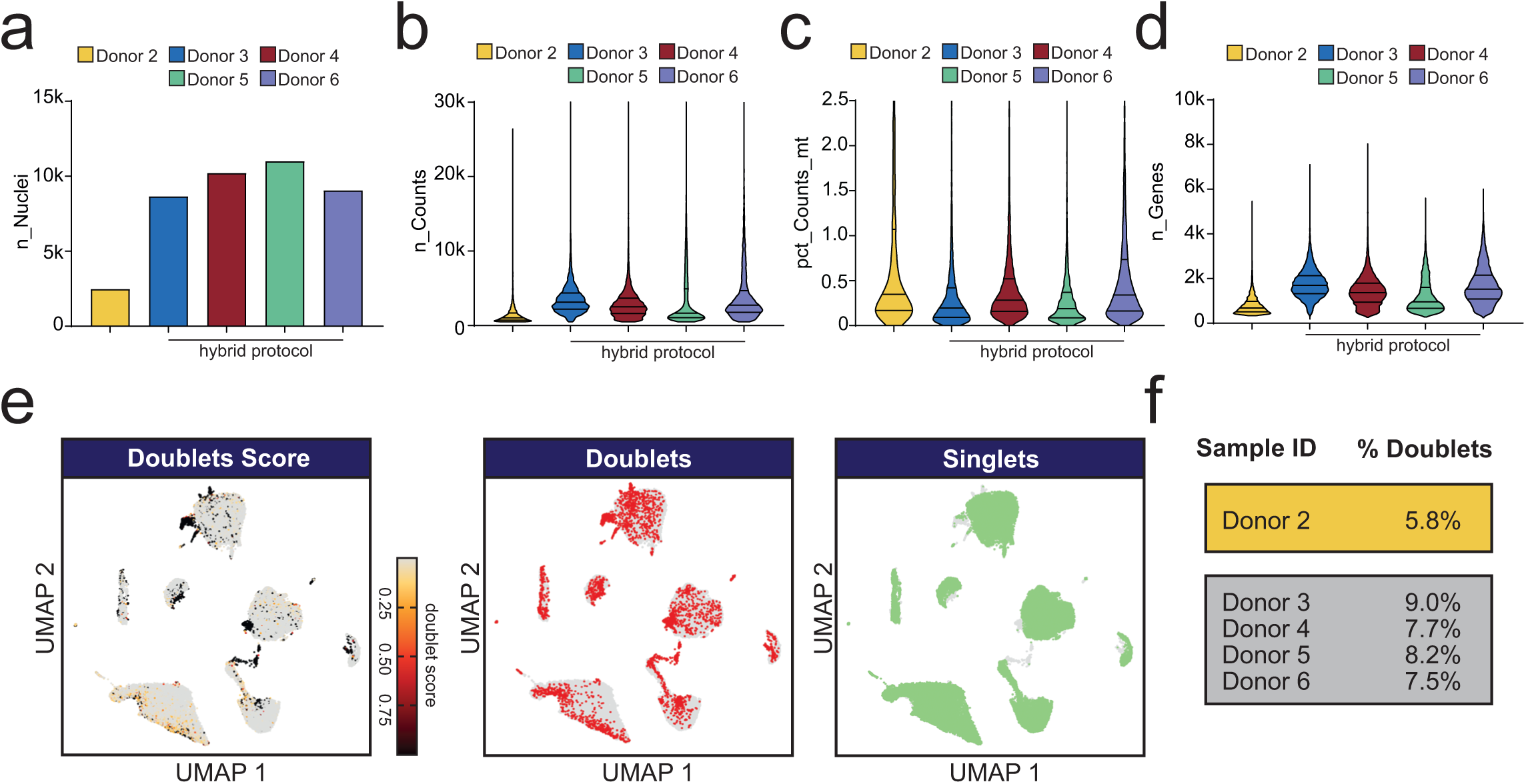
Post-sequencing quality metrics and doublet detection for nuclei isolated with two-step clean-up approach vs. hybrid method. **(a)** Bar chart showing number of nuclei recovered per sample (n_Nuclei). Violin plots showing **(b)** the median of UMIs per nucleus (n_Counts), **(c)** median percentage of reads mapping to mitochondrial RNA (pct_Counts_mt), and **(d)** median of genes detected per nucleus (n_Genes). For the violin plots, the median and interquartile range are indicated by solid lines. Samples processed using hybrid protocol are indicated. **(e)** UMAPs embeddings showing distribution of potential duplets identified using scDblFinder. Probability of datapoint to be doublets is indicated by *doublet.score*. Nuclei are annotated as doublets (red) or singles (green) based on *doublet.score*. **(f)** Percentage of doublets identified per sample. UMAPs=uniform manifold approximation and projection.

**Supplementary Figure 6.**
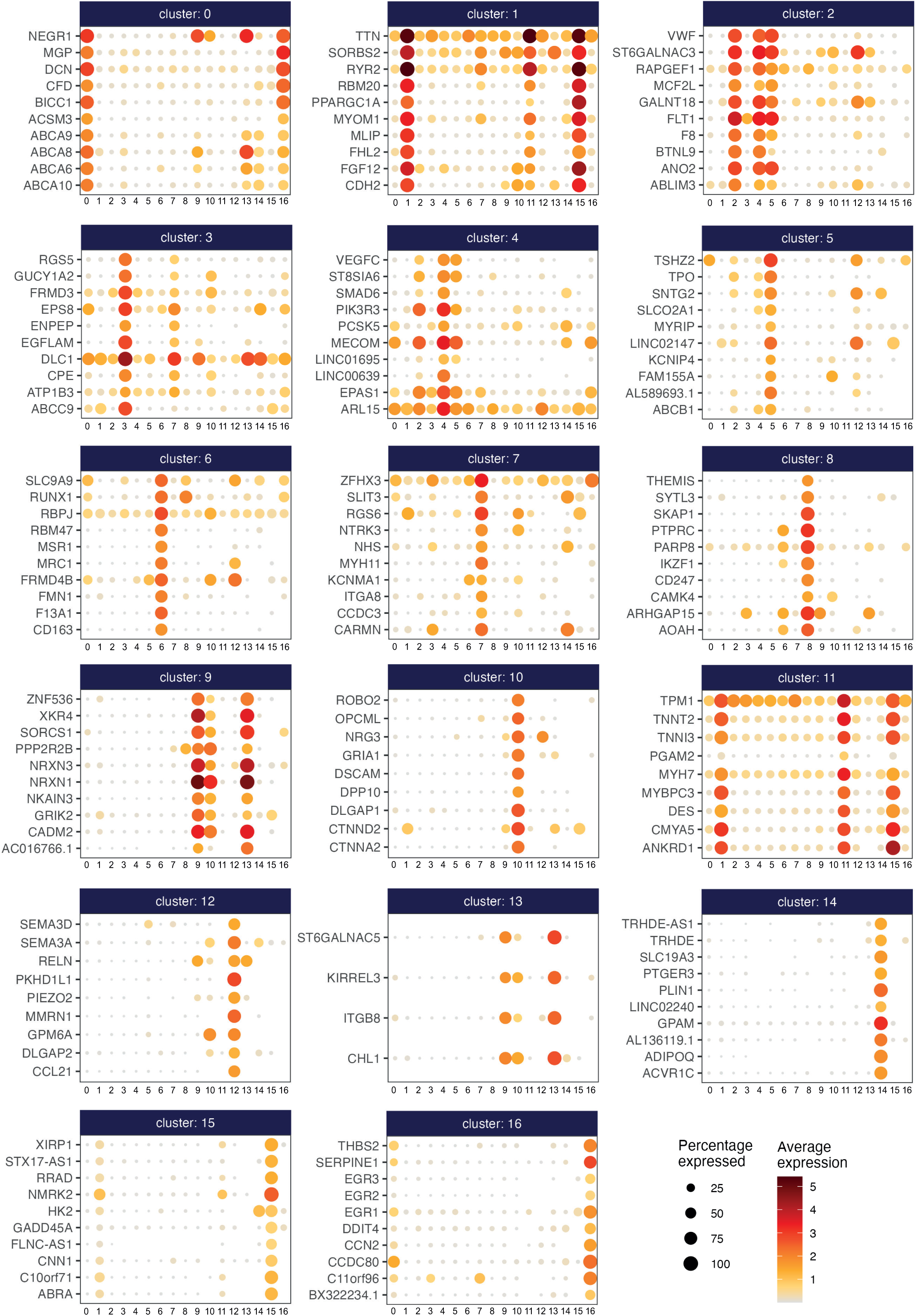
Bubble plots of top 10 differentially expressed genes per identified cluster. Bubble plots showing the top 10 differentially expressed genes (DEGs) for each of the 17 clusters identified by cluster analysis. For each gene, bubble size represents the percentage of nuclei in which the gene is differentially expressed, while bubble color indicates the average expression level within the cluster.

## REFERENCES

1. Gardner RS, Tucker NR, Amancherla K. Leveraging Single-Cell Technologies to Advance Understanding of Myocardial Disease. Circ Res. 2026;138:e326002. doi: 10.1161/circresaha.125.326002

2. Molla Desta G, Birhanu AG. Advancements in single-cell RNA sequencing and spatial transcriptomics: transforming biomedical research. Acta Biochim Pol. 2025;72:13922. doi: 10.3389/abp.2025.13922

3. Sant P, Rippe K, Mallm JP. Approaches for single-cell RNA sequencing across tissues and cell types. Transcription. 2023;14:127–145. doi: 10.1080/21541264.2023.2200721

4. Rienks M, Papageorgiou AP, Frangogiannis NG, Heymans S. Myocardial extracellular matrix: an ever-changing and diverse entity. Circ Res. 2014;114:872–888. doi: 10.1161/circresaha.114.302533

5. Park SY, Gifford JR, Andtbacka RH, Trinity JD, Hyngstrom JR, Garten RS, Diakos NA, Ives SJ, Dela F, Larsen S, et al. Cardiac, skeletal, and smooth muscle mitochondrial respiration: are all mitochondria created equal? Am J Physiol Heart Circ Physiol. 2014;307:H346–352. doi: 10.1152/ajpheart.00227.2014

6. Zhang X, Li T, Liu F, Chen Y, Yao J, Li Z, Huang Y, Wang J. Comparative Analysis of Droplet-Based Ultra-High-Throughput Single-Cell RNA-Seq Systems. Molecular Cell. 2019;73:130–142.e135. doi: 10.1016/j.molcel.2018.10.020

7. Litviňuková M, Talavera-López C, Maatz H, Reichart D, Worth CL, Lindberg EL, Kanda M, Polanski K, Heinig M, Lee M, et al. Cells of the adult human heart. Nature. 2020;588:466–472. doi: 10.1038/s41586-020-2797-4

8. Kanemaru K, Cranley J, Muraro D, Miranda AMA, Ho SY, Wilbrey-Clark A, Patrick Pett J, Polanski K, Richardson L, Litvinukova M, et al. Spatially resolved multiomics of human cardiac niches. Nature. 2023;619:801–810. doi: 10.1038/s41586-023-06311-1

9. Youness M, Ekhteraei-Tousi S, Nagaraju CK, Puertas RD, Thienpont B, Rega F, Sipido KR, Roderick HL. Location and aetiology are determinants of fibroblast activation and heterogeneity in the failing human heart. Genome Med. 2025. doi: 10.1186/s13073-025-01580-z

10. Reichart D, Lindberg EL, Maatz H, Miranda AMA, Viveiros A, Shvetsov N, Gärtner A, Nadelmann ER, Lee M, Kanemaru K, et al. Pathogenic variants damage cell composition and single cell transcription in cardiomyopathies. Science. 2022;377:eabo1984. doi: 10.1126/science.abo1984

11. Maatz H, Lindberg EL, Adami E, López-Anguita N, Perdomo-Sabogal A, Cocera Ortega L, Patone G, Reichart D, Myronova A, Schmidt S, et al. The cellular and molecular cardiac tissue responses in human inflammatory cardiomyopathies after SARS-CoV-2 infection and COVID-19 vaccination. Nat Cardiovasc Res. 2025;4:330–345. doi: 10.1038/s44161-025-00612-6

12. Selewa A, Luo K, Wasney M, Smith L, Sun X, Tang C, Eckart H, Moskowitz IP, Basu A, He X, et al. Single-cell genomics improves the discovery of risk variants and genes of atrial fibrillation. Nat Commun. 2023;14:4999. doi: 10.1038/s41467-023-40505-5

13. Eraslan G, Drokhlyansky E, Anand S, Fiskin E, Subramanian A, Slyper M, Wang J, Van Wittenberghe N, Rouhana JM, Waldman J, et al. Single-nucleus cross-tissue molecular reference maps toward understanding disease gene function. Science. 2022;376:eabl4290. doi: 10.1126/science.abl4290

14. Tran MN, Maynard KR, Spangler A, Huuki LA, Montgomery KD, Sadashivaiah V, Tippani M, Barry BK, Hancock DB, Hicks SC, et al. Single-nucleus transcriptome analysis reveals cell-type-specific molecular signatures across reward circuitry in the human brain. Neuron. 2021;109:3088–3103.e3085. doi: 10.1016/j.neuron.2021.09.001

15. Dai R, Zhang M, Chu T, Kopp R, Zhang C, Liu K, Wang Y, Wang X, Chen C, Liu C. Precision and Accuracy in Quantitative Measurement of Gene Expression from Single-cell/nucleus RNA Sequencing Data. Genomics Proteomics Bioinformatics. 2025;23. doi: 10.1093/gpbjnl/qzaf077

16. Kim N, Kang H, Jo A, Yoo SA, Lee HO. Perspectives on single-nucleus RNA sequencing in different cell types and tissues. J Pathol Transl Med. 2023;57:52–59. doi: 10.4132/jptm.2022.12.19

17. Hao Y, Hao S, Andersen-Nissen E, Mauck WM, 3rd, Zheng S, Butler A, Lee MJ, Wilk AJ, Darby C, Zager M, et al. Integrated analysis of multimodal single-cell data. Cell. 2021;184:3573–3587.e3529. doi: 10.1016/j.cell.2021.04.048

18. Germain PL, Lun A, Garcia Meixide C, Macnair W, Robinson MD. Doublet identification in single-cell sequencing data using scDblFinder. F1000Res. 2021;10:979. doi: 10.12688/f1000research.73600.2

19. Pimpalwar N, Czuba T, Smith ML, Nilsson J, Gidlöf O, Smith JG. Methods for isolation and transcriptional profiling of individual cells from the human heart. Heliyon. 2020;6:e05810. doi: 10.1016/j.heliyon.2020.e05810

20. Larson A, Chin MT. A method for cryopreservation and single nucleus RNA-sequencing of normal adult human interventricular septum heart tissue reveals cellular diversity and function. BMC Med Genomics. 2021;14:161. doi: 10.1186/s12920-021-01011-z

21. Safabakhsh S, Sar F, Martelotto L, Haegert A, Singhera G, Hanson P, Parker J, Collins C, Rohani L, Laksman Z. Isolating Nuclei From Frozen Human Heart Tissue for Single-Nucleus RNA Sequencing. Curr Protoc. 2022;2:e480. doi: 10.1002/cpz1.480

22. Araten S, Mathieu R, Jetly A, Shin H, Hilal N, Zhang B, Morillo K, Nandan D, Sivankutty I, Chen MH, et al. High-quality nuclei isolation from postmortem human heart muscle tissues for single-cell studies. J Mol Cell Cardiol. 2023;179:7–17. doi: 10.1016/j.yjmcc.2023.03.010

